# Line-FRAP, a versatile method based on fluorescence recovery after photobleaching to measure diffusion rates *in vitro* and *in vivo*

**DOI:** 10.1101/2020.07.01.181750

**Authors:** Debabrata Dey, Shir Marciano, Gideon Schreiber

**Affiliations:** Department of Biomolecular Sciences, Weizmann Institute of Science

## Abstract

A cell is a densely packed conglomerate of macromolecules, where diffusion is essential for their function. The crowded conditions may affect diffusion both through hard (occluded space) and soft (weak, non-specific) interactions. Multiple-methods have been developed to measure diffusion rates at physiological protein concentrations within cells, however, each of them has its limitations. Here, we introduce Line-FRAP, a method based on measuring recovery of photobleaching under a confocal microscope that allows diffusion rate measurements for fast diffusing molecules to be measured in versatile environments using standard equipment. Implementation of Line mode to the classical FRAP technique greatly improves the time resolution in data acquisition, from 20-50 Hz in the classical mode to 800 Hz in the line mode. We also introduce an updated method for data analysis to obtain diffusion coefficients in various environments, with the number of pixels bleached at the first frame after bleaching being a critical parameter. We evaluated the method using different proteins either chemically labelled or by fusion to YFP. The calculated diffusion rates were comparable to literature data as measured *in vitro*, in HeLa cells and in *E.coli.* Diffusion coefficients in HeLa was ~2.5-fold slower and in *E. coli* 15-fold slower than measured in buffer. Moreover, we show that increasing the osmotic pressure on *E.coli* further decreases diffusion, till a point where proteins stop to move. The method presented here is easy to apply on a standard confocal microscope, fits a large range of molecules with different sizes and provides robust results in any conceivable environment and protein concentration for fast diffusing molecules.

## Introduction

For cells to function, molecules of different sizes and composition diffuse in the crowded environment to reach their targets(Dix & Verkman, 2008; Goodsell, 1991; Verkman, 2002). Fick’s law, which predicts Diffusion driven by Brownian motion, is strictly valid for ideal solutions (with infinitely dilute concentrations of the solutes). It overlooks interactions between the moving species and with the surrounding (Agutter et al., 2000; Haugh, 2009; Saxton, 2012). Simulations and experiments describing molecular motions in a complex environment is a stiff challenge due to the various soft and hard interactions among the moving species(Baumann et al., 2010; Lenzi et al., 2016; Schavemaker et al., 2018). The classical theory of Stokes-Einstein regarding particle diffusion in ideal solution states that the diffusion constant of a particle *A* (*D_A_*) is a function of its dimensions and the solution viscosity(Chandrasekhar, 1943; Einstein, 1905; Sutherland, 1905).

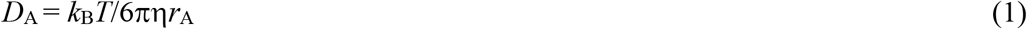

where η is the solvent viscosity, *k_B_T* is the product of the Boltzmann constant and the absolute temperature, *r_A_* is the hydrodynamic radius of the diffusing particle and *D_A_* is a measure of the mean squared displacements per unit time.(Sutherland, 1905) Interactions with other molecules that lower the mobility will be reflected by lower *D_A_* values. Alternatively, if the motion of the molecule is increased because of the assistance from another faster moving dynamic molecule/surrounding environments, higher *D_A_* values will be recorded. Both of these are known as anomalous diffusion(Haugh, 2009; Saxton, 2012). Accurate measurements of *D_A_* values in the complex environment should correlate with the effective hydrodynamic radius, which takes into account the interactions the molecule is engaged in(Day & Kenworthy, 2009; He & Niemeyer, 2003; Mueller et al., 2010).

Over the years a wide range of different techniques have been developed to measure diffusion of different species inside the living cell. Most of these are fluorescent based. A comprehensive detail of different existing fluorescence based microscopy techniques and their utility/limitations are shown in supporting information, Table S1. Among them, Fluorescence recovery after Photobleaching (FRAP) (Kitamura et al., 2015; Lippincott-Schwartz et al., 2003; Phair & Misteli, 2000; Verkman, 2003), fluorescence correlation microscopy (FCS) (Bacia et al., 2006; Cluzel et al., 2000; Dauty & Verkman, 2004; Ramadurai et al., 2009) and single particle tracking (SPT) (Manley et al., 2008) are the most popular. On the other hand, techniques like RCIS (Digman & Gratton, 2009), FSM (Waterman-Storer et al., 1998) or PIPE (Gura Sadovsky et al., 2017) are less frequently used. These techniques, separately or in combination, give reasonable diffusion values within the accepted ranges reported in the literature(Mika & Poolman, 2011; Miyawaki, 2011). Still, diffusion values reported by different research groups vary, predominantly inside the complex environment of the cell(Kingsley et al., 2018). In addition, the assumptions of different mathematical models and imposed boundary conditions on the data processing of the different techniques can influence the results, which ultimately leads to a diversified range of diffusion values(Kingsley et al., 2018). Even for a specific molecule of interest using a single instrumental technique, diversified diffusion values create confusion and misunderstanding about the model and its underlying assumptions(Kingsley et al., 2018).

FCS is considered the gold standard for calculating diffusion coefficients in solution phase. However, it requires the measured particles to be very dilute (as low as ~1 - 10 nM), and a high quantum yield of the measured particle. This imposes sever limitations for measurements in crowded environments like inside the living cell. Moreover, if molecules have some bound fraction that is immobile, FCS remains completely blind to this fraction(Ries & Schwille, 2008; Wachsmuth, 2014). In addition, FCS has limitations regarding accurate detection of slowly binding molecules(Wachsmuth, 2014).

FRAP, is the most widely used technique by experimental biologists because of the relative ease of performing confocal FRAP in the majority of the biological domains. This technique is also very sensitive towards the bound or immobile fractions of the molecules under study(Costantini & Snapp, 2013; González-González et al., 2012; Hardy, 2012; Zuleger et al., 2013). However, there are major limitation associated with the FRAP technique. For one is the relative slow data accumulation rate, making it insensitive to fast diffusing particles(Wachsmuth, 2014). Second, the conversion of FRAP rates to *D_A_* is problematic, and requires making multiple assumptions(Angelides et al., 1988; Axelrod et al., 1976; Braga et al., 2004; Deschout et al., 2010; Kang et al., 2009; Seiffert & Oppermann, 2005; Smisdom et al., 2011; Soumpasis, 1983; Sprague et al., 2004; Waharte et al., 2010). Here, we present a modified FRAP method to calculate *D_A_* from *in vitro* to *in vivo* and provide robust diffusion data, similar to those determined using FCS. To overcome the slow data acquisition of the confocal microscope we devised a line-FRAP method, where a single scan along a line is repeated at ~1000 Hz. The use of two simultaneous scanners reduced the dead time of the instrument, resulting in further increase in efficiency of the line FRAP(Braeckmans et al., 2007). Careful evaluation of the size of the bleached area resulted in robust diffusion results for proteins and small molecules in varied conditions, including simple buffers, eukaryotic cells and the dense milieu of the bacterial cytoplasm under varying osmotic stresses.

## Results

### Theoretical background on extracting diffusing rates from FRAP data

Extracting diffusion coefficient values from FRAP recovery profiles is not straightforward(Angelides et al., 1988; Axelrod et al., 1976; Braga et al., 2004; Deschout et al., 2010; Kang et al., 2009; Seiffert & Oppermann, 2005; Smisdom et al., 2011; Soumpasis, 1983; Sprague et al., 2004; Waharte et al., 2010). The easiest way is to calculate half time of recovery (τ1/2) of the FRAP spot, which is defined as the time required for 50% recovery of the bleach spot(Lajoie et al., 2007; Salmon et al., 1984; Stricker et al., 2002; Zotter et al., 2017). Nevertheless, as also shown previously, τ1/2 measurements may actually be misleading, particularly as they do not take into account the processes occurring during bleach(Axelrod et al., 1976; Soumpasis, 1983). The classic equation derived by Soumpasis(Soumpasis, 1983) relating FRAP to the diffusion rate is:

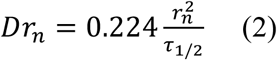

This equation assumes a uniform radius of the bleach area (r_n_)(Axelrod et al., 1976; Soumpasis, 1983). This relation is strictly valid for a pure isotropic diffusion model(Axelrod et al., 1976; Soumpasis, 1983). However, the main assumption i.e., diffusion during photobleaching is neglected, making its use for real FRAP experimental conditions limited. For normal laser scanning confocal microscopes, the relatively longer scanning durations increases the probability of significant amount of diffusions even during the photobleaching time pulse. This phenomenon obviously will be more common for faster diffusing molecules. Therefore, *D* values calculated from equation 2 can lead to underestimation of actual *D* in many systems(Braga et al., 2004; Kang et al., 2009; Pucadyil & Chattopadhyay, 2006; Weiss, 2004). Kang et.al. (Kang et al., 2012) solved these limitations by incorporating a new parameter, the effective bleach radius r_e_. The final usable form of the modified equation is:

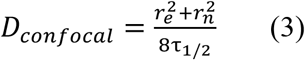

When bleaching is instantaneous (i.e. for very short time pulse laser, r_e_ = r_n_) 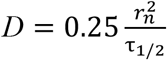, which is similar to equation 2 (Kang et al., 2012). For the common case, where r_e_ ≠ r_n_ we determined experimentally r_e_, r_n_ and τ_1/2_ in different conditions for line-FRAP, averaging 30 measurements for each sample and then using the effective bleach radius r_e_ in equation 3 to determine *D*_confocal_.

### Line FRAP methodology

Fig. 1A shows a pictorial representation of the Line-FRAP method performed in a life HeLa cell expressing CFP (Cyan Fluorescent protein), showing both the fluorescence and transmission Channels. The small red circle denotes the bleaching area, which is done by a Clip tornado (of 4-pixel size, 1 pixel length equals to 0.207 micro meter) as defined by the Olympus confocal system. The white line is the scanning line, which is performed at a rate of 800 Hz, to follow FRAP as a function of time. As we record only a line, the method is named as Line-FRAP (Braeckmans et al., 2007). Figure 1B shows the Line-FRAP cross section profile as a function of time. The vertical direction of the line denotes the time progress. The vertical red lines in the left panel shows the Region of Interest (ROI), from which the fluorescence intensity over time is plotted (Figure 1C). The horizontal rectangular area marked as 1 (red color) in Fig. 1B right panel denotes the time frame immediate after the photo-bleach pulse (the highly intense green is the bleach pulse). The time interval between each line measurement is 1.25 millisecond. The darker region reflects the bleached area. A typical example of a FRAP recovery profile (averaged over 30 individual shots) is shown in Figure 1C. The red line denotes fitting the data to a double exponential progression curve.

**Figure 1:**
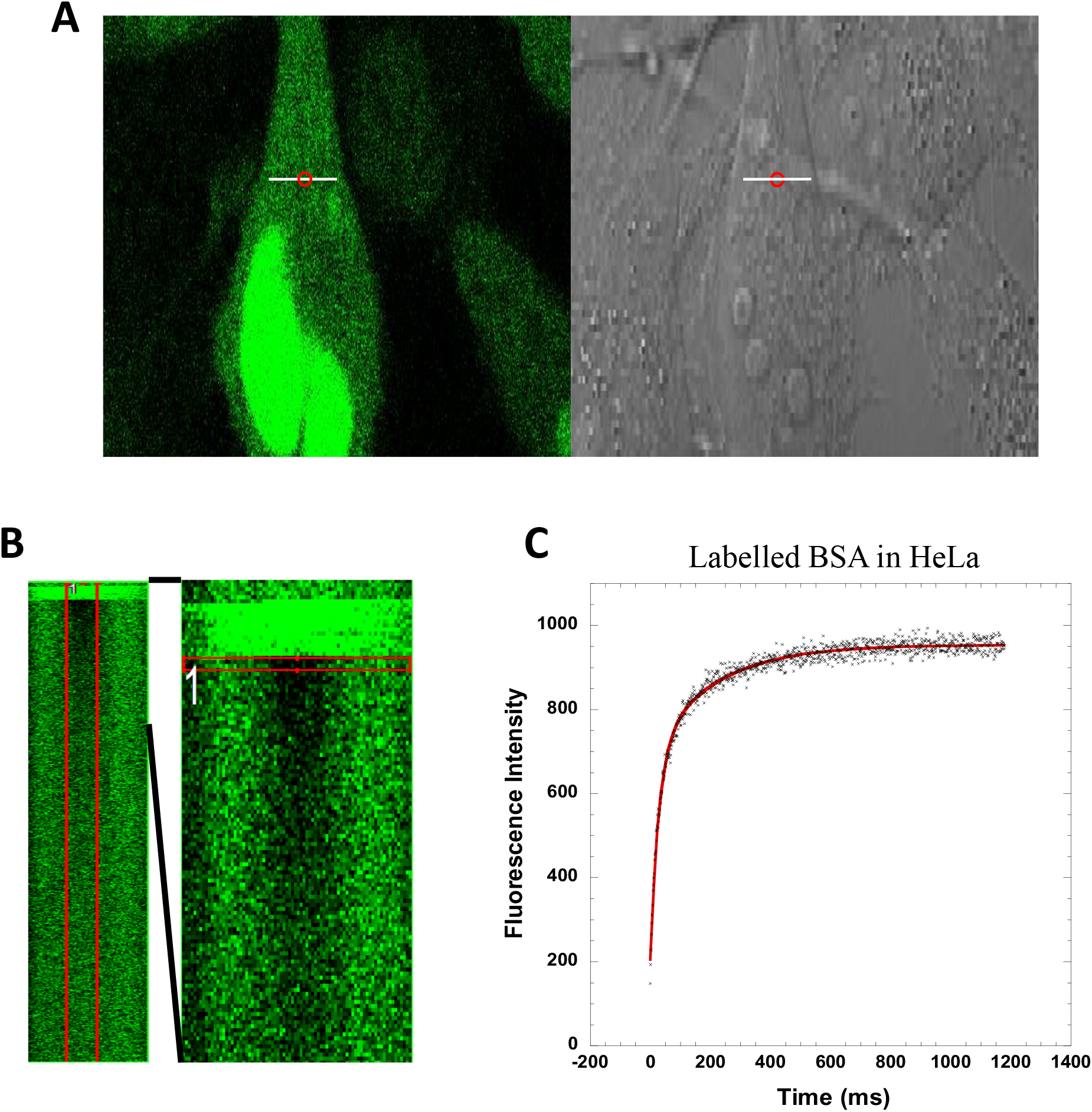
(A) Line-FRAP performed in a CFP transfected HeLa cell. White line represents the scanning line; Red circular area denotes the bleaching area. (B): A single scanning line as a function of time (in vertical direction) including the bleach. The red rectangular marks the region of interest (ROI), which was used for analysis. A zoomed view of Line-FRAP profile, is also shown where horizontal rectangular area (red colored marks ROI) denotes the fluorescence intensity as a function of length across the scanning line. (C): Average of 30 FRAP recovery curves as a function of time. (N=30; Correlation coefficient R=0.98)

Several mathematical functions were tested to fit the FRAP recovery data. The bi-exponential function gave a better fit than a single exponent function, not only in complex environments but also in simple PBS buffer solution. The two-rate constant parameter *k*_1_ and *k*_2_ with their associated amplitudes (a_1_ and a_2_) were used to calculate τ_1/2_ using equation 4.

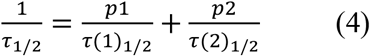

Where p_1_ and p_2_ are the fraction amplitudes of *k*_1_ and *k*_2_ and τ(1)_1/2_ τ(2)_1/2_ are the respective half-recovery times. The reasons we observe more than one exponent in the recovery curve is the diffusive boundary between the bleached area and its surrounding (Fig. 2). This relates both to dispersion of the laser light and to diffusion occurring during the bleach time.

**Figure 2:**
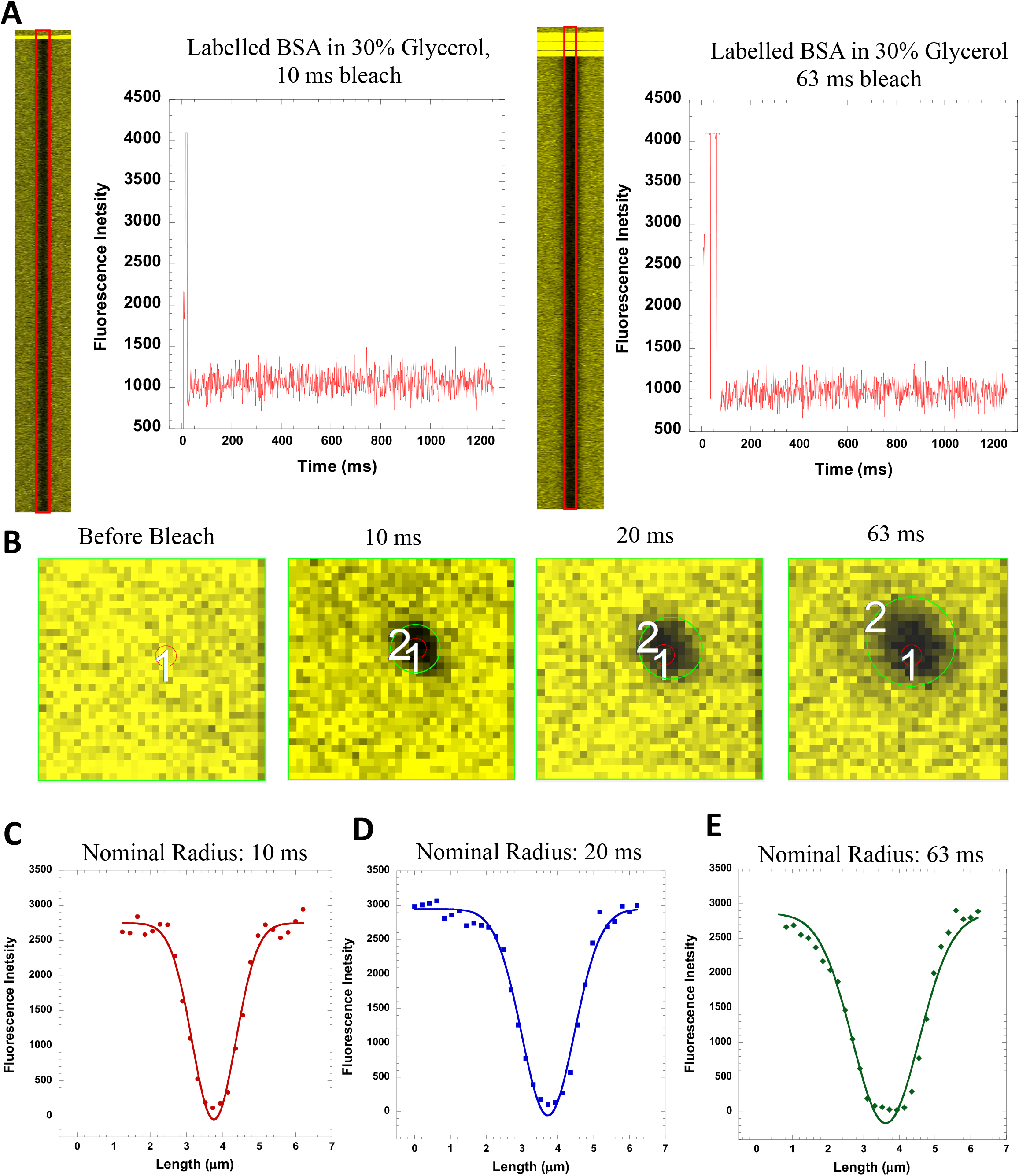
(A) Line FRAP profile of a precipitated (fixed) sample (BSA labeled with CF514 Dye in 30% Glycerol in PBS buffer) after 10 ms and 63 ms bleach pulse. Corresponding recovery profiles over time are shown. (B): Nominal bleach areas (before and after the bleach) for the precipitated samples. The green circular area marked 2, denotes the nominal bleach done by the laser when 10, 20 and 63 ms bleach pulses were used and the red circular area 1 denotes the bleach area defined by the clip tornado of 4 pixel (Inherent laser settings defined by Olympus confocal system). (C-E): Determination of nominal bleach radius (rn) from the fluorescence intensity profiles for (C) 10, (D) 20 and (E) 63 ms bleach pulse fitted using a Gaussian equation. (N=20; R=0.99 for each of the fits)

### Determination of r_n_ and r_e_ of the Bleach area

Equation 3 is very sensitive to r_n_ and r_e_ values, which are squared, while τ_1/2_ is not. Kang et.al (Kang et al., 2012) estimated r in a slow diffusing system that allows 50 – 100 ms time lapses between frames. This is not relevant for fast diffusing systems, where recovery from bleach will occur within the first few frames. To obtain r_n_ values that are disconnected from diffusion we prepared a fluorescent protein sample that forms an aggregate. The sample consisted of BSA labelled with CF514 dye in PBS buffer and 30 percent (v/v) glycerol. Figure 2A shows Line FRAP profiles with corresponding fluorescence recovery over time traces for this sample. No FRAP were observed after photo bleach pulses of 10 ms and 63 ms. Figure 2B shows the bleach radius after 10, 20 and 63 ms of bleach pulses. The red circle is the set bleach of the laser (4 pixels). As seen, the size of the bleached area increases with increasing bleach times, which relates to the diffraction of the laser light, with the laser power set to 100% intensity in all cases(Kitamura & Kinjo, 2018). To quantify the signal, we plotted the intensity versus the position along the horizontal scanning line (Figure 2 C-E). To improve the signal to noise, we averaged 30 individual measurements and fitted them to a Gaussian (equation 5):

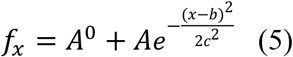

where *A*^0^ is the offset, *A* the amplitude, *b* the midpoint and *c* the width, which is related to the width at half maximum (FWHM) by FWHM = 2(2ln2)^1/2^*c*. Figure 2 (C-E) shows the Gaussian fits for post-bleach fluorescence profiles for 10, 20 and 63 ms of bleach pulses respectively.

Nominal radius values are assumed to be fixed for a given microscope setting and laser. Not so for the effective radius r_e_, which depends also on the diffusion during bleach. Therefore it is mandatory to measure the effective bleach radius for each sample separately. Figure 3A shows Line-FRAP progression profiles and curves of BSA labelled with CF514 in PBS buffer, using 10 and 63 ms bleach pulse times. Figure 3B shows the fluorescence intensity along the horizontal line (after the photo bleach in Line-FRAP mode) for a single measurement. Figure 3C is an average of 30 measurements and figure 3D shows a gaussian fit of these data. Ambiguity remains over the percent recovery to be used for calculating r_n_ and r_e_ values. Kang et.al used 90% fluorescence signal recovery for r_e_ as the diameter of the effective bleach circle (Kang et al., 2012). However, this is an arbitrary choice, and therefore we preferred to obtain this value from equation 3 by comparing FRAP diffusion rates to those obtained by FCS for the same molecule. For this purpose we chose BSA, which diffusion rate has been estimated to be 63 and 51 μm^2^s^−1^ using light scattering(Gaigalas et al., 1992), and FCS(Krouglova et al., 2004) measurements at 25°C. Using the fitted *c* parameter in equation 5, we found that r_n_ and r_e_ should be taken at 60% of total fluorescence signal recovery, which is equal to 1.38\**c*. Accordingly, r_n_ values for 10, 20 and 63 ms bleach pulses calculated from the Gaussian fits are 0.745, 0.98 and 1.27 μm. These values show perfect agreement with the number of dark pixels present in the Figure 2B. Table 1 shows the parameters used to obtain diffusion rates for BSA after 10 and 63 ms bleach times. While the bleach recovery rates and r_e_ are very different for 10 and 63 ms bleach time(Kitamura & Kinjo, 2018), the calculated diffusion coefficient is within 10% from each other, and fit previous measurements(Gaigalas et al., 1992), (Krouglova et al., 2004). Another outcome from this calibration experiment is that the calculation procedure remained exactly the same, independent on whether 10 or 63 ms bleach time were used, despite the very large differences in all measured parameters.

**Figure 3:**
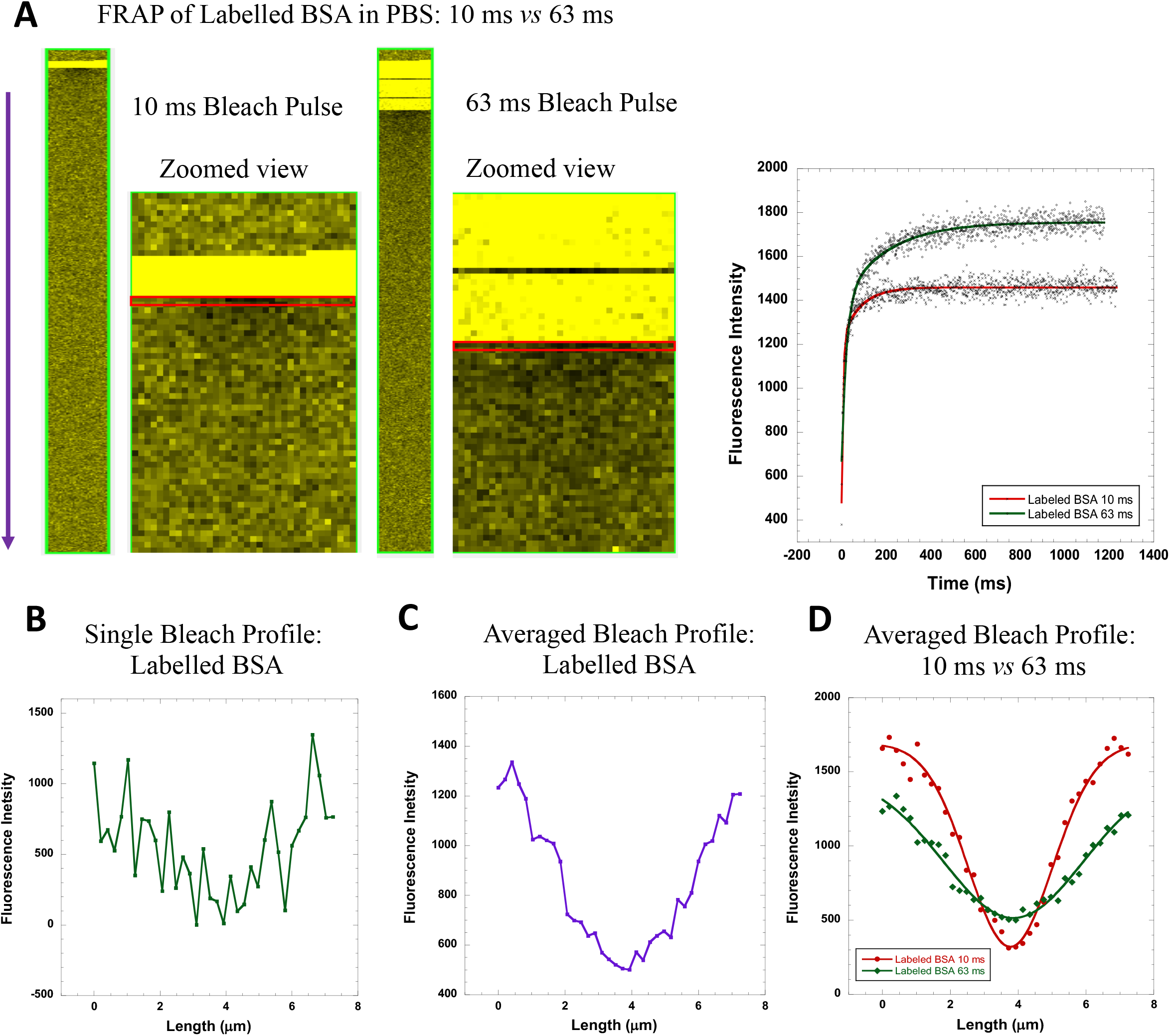
(A) Fluorescence recovery (FRAP) after photobleaching of BSA labelled with CF514 in PBS. Comparative data of 10 and 63 ms bleach pulse fitted to double exponential equation are shown. (N=30; R=0.91 for 10ms and 0.97 for 63ms fit) (B-D): Determination of effective bleach sizes from the fluorescence intensity profiles for 10 and 63 ms bleach pulse. (B) Single run data, (C) average of 30 measurements. (D) Gaussian fits for the average bleach sizes after 10 and 63 ms bleach pulse. (N=30; R= 0.99 for both of the fits)

**Table 1:** Diffusion Coefficients calculated from FRAP data for labeled BSA in PBS buffer at different bleach time durations

| Bleach Pulse duration | $k_1$<br>$\text{ms}^{-1}$ | $k_2$<br>$\text{ms}^{-1}$ | $a_1$ | $a_2$ | $p_1$ | $p_2$ | $\tau_1$ | $\tau_2$ | $\tau_{1/2}$<br>(ave) | $c_n$<br>$\mu\text{m}$ | $r_n$<br>$\mu\text{m}$ | $c_e$<br>$\mu\text{m}$ | $r_e$<br>$\mu\text{m}$ | $D_{\text{confocal}}$<br>$\mu\text{m}^2 \text{s}^{-1}$ |
| --- | --- | --- | --- | --- | --- | --- | --- | --- | --- | --- | --- | --- | --- | --- |
| 10ms | 0.16 | 0.012 | 736 | 241 | 0.75 | 0.25 | 0.004 | 0.058 | 0.0056 | 0.54 | 0.74 | 1.15 | 1.59 | 68.5 |
| 10ms | 0.16 | 0.013 | 1125 | 307 | 0.78 | 0.21 | 0.004 | 0.053 | 0.0054 | 0.54 | 0.74 | 1.15 | 1.59 | 71.2 |
| 63ms | 0.046 | 0.005 | 726 | 359 | 0.67 | 0.33 | 0.015 | 0.14 | 0.0214 | 0.92 | 1.27 | 2.05 | 2.83 | 54.9 |
| 63ms | 0.044 | 0.005 | 770 | 373 | 0.67 | 0.33 | 0.016 | 0.14 | 0.0222 | 0.92 | 1.27 | 2.05 | 2.83 | 54.2 |
| 63ms | 0.046 | 0.005 | 466 | 254 | 0.65 | 0.35 | 0.015 | 0.13 | 0.0219 | 0.92 | 1.27 | 2.05 | 2.83 | 56.3 |

The rate constants ( $k$ ) and amplitudes ( $a$ ) are from the double-exponential fit of the FRAP data. The $c$ values were obtained from Gaussian fits using equation 5. The respective radius ( $r$ ) are calculated assuming the respective bleach circles diameter lies at 60% height from the bottom of the Gaussian Profiles ( $r=1.38*c$ ).

### Comparing Line FRAP to standard FRAP

The main advantage of Line FRAP over regular FRAP is the much improved time resolution. In the Supporting Information (Figures S1 to S3) we show FRAP data, using the same microscope setup and CF514 labelled BSA as used in Figure 3, but applying classical XY FRAP. While using Line FRAP data acquisition was at 800 Hz, using standard FRAP it was 8.5 and 17 Hz for squares of 72×72 (14.90 μm x14.90 μm) or 36×36 pixels (7.45 μm x7.45 μm) respectively (Figures S1A and B show the first frame after bleach). As can be seen, 36×36 pixels is the smallest square possible to see the boundaries of the bleach spot. In Figure S1C we show XY FRAP after 10 ms bleach pulse, which obviously results in poor bleach in the first frame (58 ms after bleach). For comparison, we show the time dependent recovery and the first frames after bleach for a 10 ms bleach pulse in Figure S1D. From Figure S1 (A-D), It is clear that Line FRAP (time frame resolution of ~1.25 ms as a line) provides far superior information compared to the existing classical FRAP techniques. In Figures S2 and S3 we show an analysis of the standard FRAP data using 63ms (Figure S2A and B) and 10 ms (Figure S2C) pulse using either a 72 (A) or 36 (B) pixel frame. For 10 ms bleach pulse only the 36 pixel frame is shown. Panel D shows for comparison the Line FRAP data for a 10 ms pulse. In the right panel we show a fit of an average of 20 acquisitions to a double-exponential equation. Figure S3 shows the bleach width for the measurements shown in Figure S2, fitted to a gaussian equation. The calculated τ_1/2_, r_n_ and r_e_ values as well as the diffusion coefficients are given in Table S2. All the calculations were done as explained for Table 1. The calculated τ_1/2_, r_n_ and r_e_ values differed between the three independent FRAP measurements using the square area method as presented in Table S2, and again between later and those obtained using the Line FRAP method. The square FRAP gave *D*_confocal_ values of 29, 90 and 21 μm^2^s^−1^, while the Line FRAP gave a value of 55 for 63 ms pulse bleach and 70 for 10 ms pulse bleach. The reason for the missing accuracy for the standard FRAP method is the long time it takes for the first frame to be recorded (during which diffusion is occurring) and the few points to fit as τ_1/2_, due to the fast diffusion and slow frame rate. Overall, this shows the advantage of the Line FRAP method over conventional FRAP for fast diffusing molecules, including proteins like BSA (MW 66.5 kDa).

### Measuring BSA diffusion in crowded environments

A rigorous test for the Line FRAP method is to retrieve diffusion rates in crowded environments. PEG8000 is a synthetic polymer, popular for its use as crowder in solution phase. Labelled BSA was added to 0, 5, 10, 15 and 20% (w/v) of PEG8000 in PBS buffer solutions. The measurements were done after 10 and 63 ms bleach pulses. Figures 4 shows FRAP recovery and post-bleach intensities for 10 ms (Figure 4A and 4B) and 63 ms (4C and 4D) bleach pulse durations. For r_n_ we used 0.745, 1.27 μm values for 10 and 63 ms bleach pulses respectively. Values of r_e_ were calculated from Figure 4B and 4D using equation 5 as detailed above. Values of τ_1/2_ were calculated using equation 4. The calculated diffusion coefficients were plotted versus the relative solution viscosity (Figure 4E). The different fitted parameters are provided in Table S3. As expected, the measured diffusion coefficients for BSA in crowded environments showed a gradual decrease with increasing relative viscosity of the PEG8000 solution(Kuttner et al., 2005). The measured diffusion values were (within the error) the same, independent on whether 10 or 63 ms bleach times were used, despite the very different recovery profiles. The results here show that Line-FRAP is capable for measuring diffusion rates in different solutions with high reproducibility.

**Figure 4:**
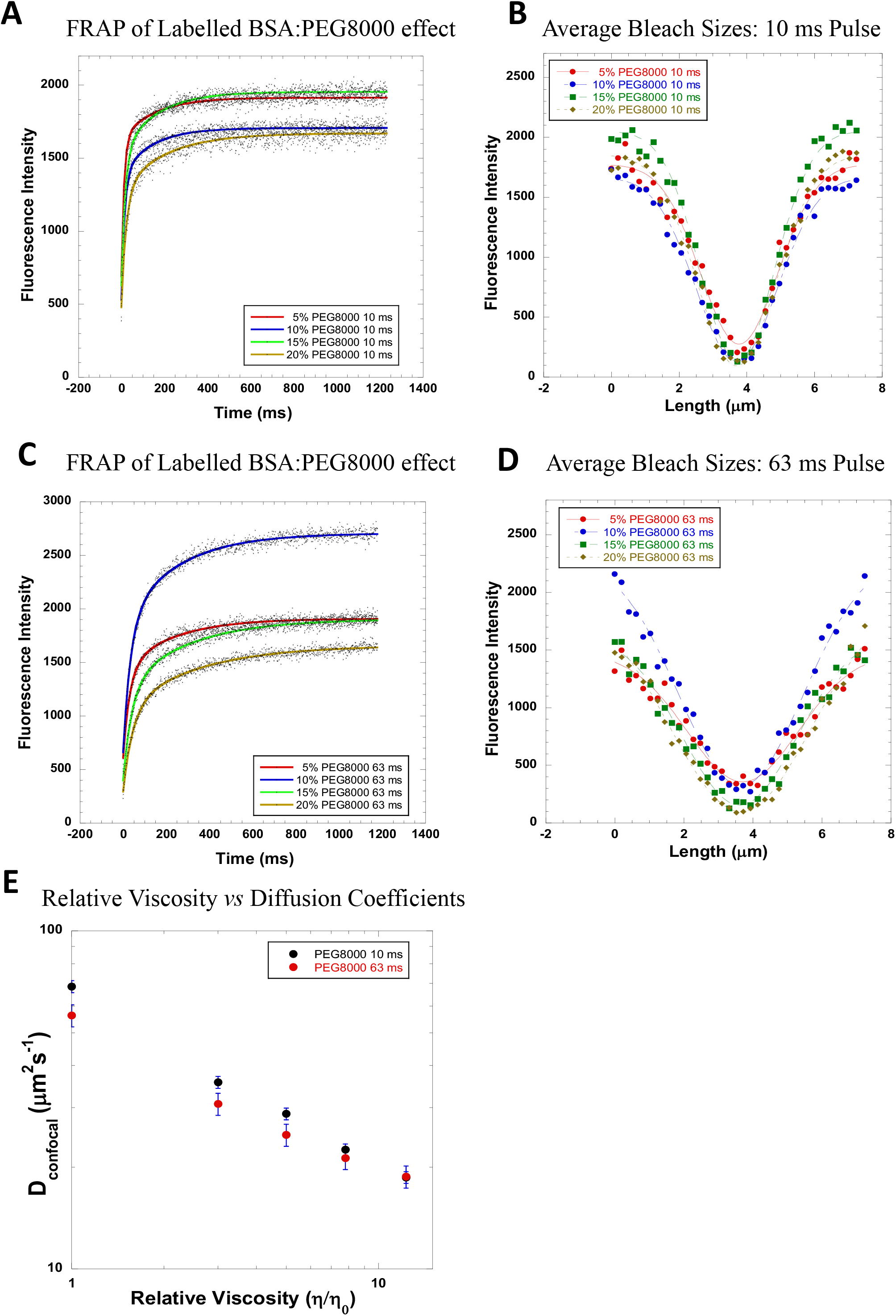
(A,C) FRAP recovery profiles for 10 and 63 ms bleach pulses of labeled BSA in PBS buffer in the presence of 5, 10, 15 and 20% PEG8000. (N=30 for all; R= 0.96 for 5, 0.97 for 10, 0.98 for 15 and 20% fits in A) (R= 0.98 for 5%, 0.99 for rest of the fits in C) (B, D) Average post bleach size of fluorescence intensity profiles after 10 ms (N=30 for all; R= 0.99 for each of the fits in B) and 63 ms bleach pulses fitted to a Gaussian equation. (N=30 for all; R= 0.98 for 5% and 0.99 for rest of the fits in D) (E): Diffusion Coefficients (in μm2s-1) as a function of Relative viscosity (η0 of water considered as 1, data are from panels A-D, final error bars are calculated after combining all the individual errors from the respective average fits).

### Diffusion Coefficient Measurements *in vivo*

Our ultimate goal is to measure Diffusion Coefficients of proteins in the crowded milieu of the cell. *In vivo* measurements introduce major technical challenges, including the variability between individual living cells. Therefore, single-cell measurements should be performed in sufficient numbers to estimate the diversity in terms of environments as well as sample variations. Here, we used two bacterial proteins from *E.coli* origin, E-fts (PDB ID **1EFU**) and baeR (PDB ID **4B09**) for our experiments. According to swiss model(Bertoni et al., 2017; Waterhouse et al., 2018), E-fts and baeR are homodimers (dimer MW = 60.36 kDa and 55.2 kDa respectively). E-fts and baeR were labelled using two methods. One was with the CF514 fluorescent dye and the second by fusing the proteins to YFP (monomer MW, 26.4 kDa). We have used the YFP protein variant with the A206K mutation to inhibit further oligomerization of the YFP(Von Stetten et al., 2012). FRAP rates were first determined in PBS (Figure 5), from which their respective diffusion coefficients were calculated as detailed above for BSA. The diffusion coefficients of E-fts and baeR labelled with CF514 were 37.2 and 46 μm^2^s^−1^ respectively. For the YFP fused proteins (with an estimated MW of 123 and 118 kDa) the diffusion coefficients were as expected slower, 22.9 and 31.4 μm^2^s^−1^ respectively (Table S4).

**Figure 5:**
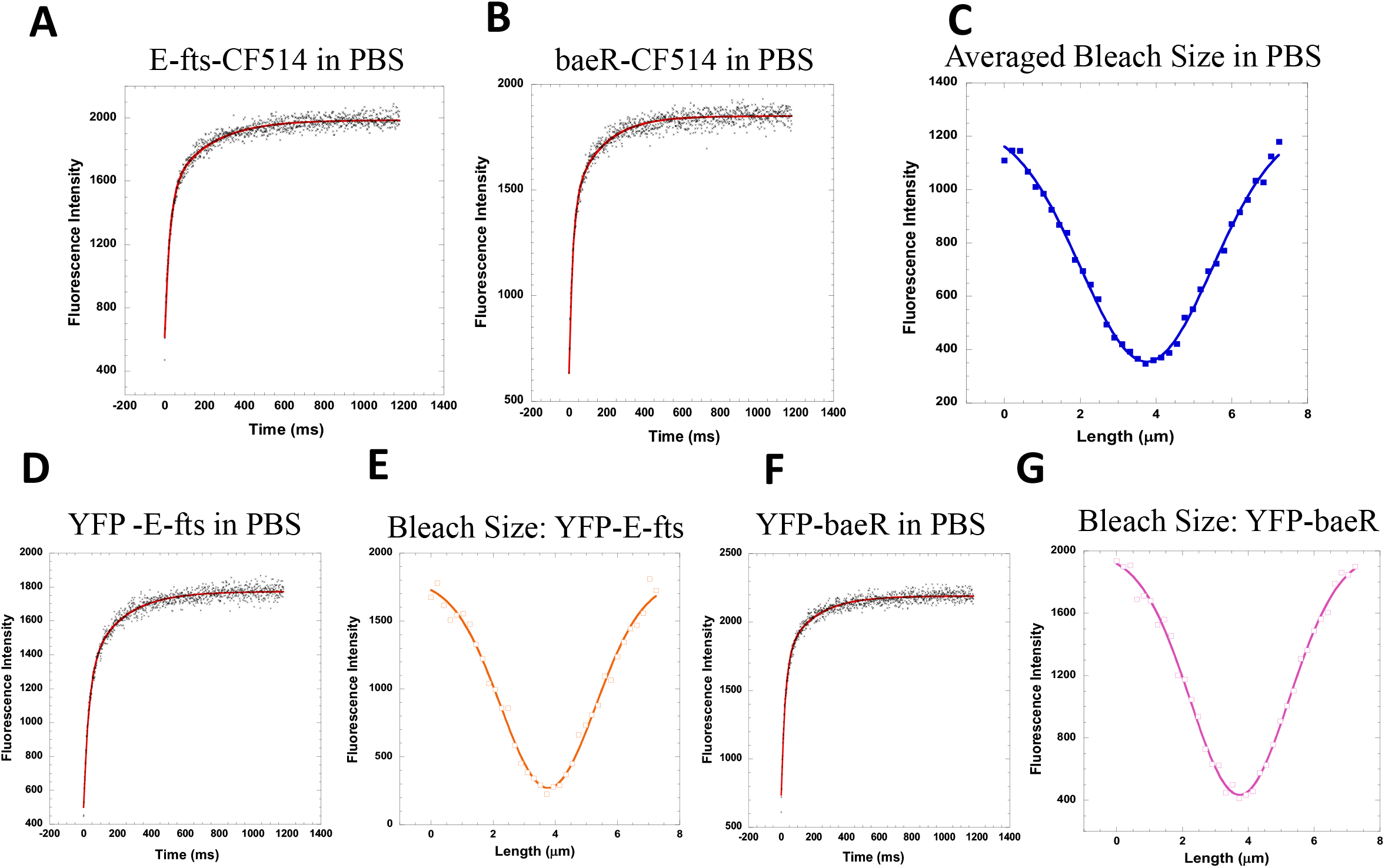
(A-C) FRAP recovery profiles of (A) E-fts (N=25; R=0.98) (B) baeR proteins (N=30; R=0.98) labeled with CF514 dye in PBS buffer. (C) Corresponding averaged post bleach size profiles for E-fts and baeR proteins fitted to a Gaussian equation (N=100; R=0.99). (D-G): FRAP recovery profiles of (D) E-fts fused with YFP (N=30; R=0.99) (F) baeR fused with YFP (N=30; R=0.99) in PBS buffer solution. Gaussian fits of averaged post bleach size profiles for (E) E-fts fused with YFP (N=30; R=0.99) (G) baeR fused with YFP (N=30; R=0.99).

Next, we used these proteins to determine their diffusion coefficients in both Eukaryotic and prokaryotic cells, as models for *in vivo* measurements. It is known that Eukaryotic cell cytoplasm are less dense than prokaryotic cell cytoplasm, which should be reflected in the diffusion coefficient(Dix & Verkman, 2008; Mika & Poolman, 2011). The 4 different proteins (E-fts and baeR labelled with CF514 or fused to YFP) were microinjected inside HeLa cells cytoplasm. Line-FRAP experiments were performed inside intact living cells (Figure. 6). FRAP rates and r_e_ values were determined, from which the diffusion coefficients were calculated (Table S4). As expected from the increased crowding of the cytoplasm of HeLa cells, slower diffusion coefficients were determined(Dix & Verkman, 2008). For CF514 labelled proteins diffusion coefficients of 19.5 and 19 μm^2^s^−1^ were determined for E-fts and baeR respectively (Table S4). Diffusion coefficients of 9.0 and 12.5 μm^2^s^−1^ were determined for these proteins fused to YFP (which results in approximately doubling the MW).

**Figure 6:**
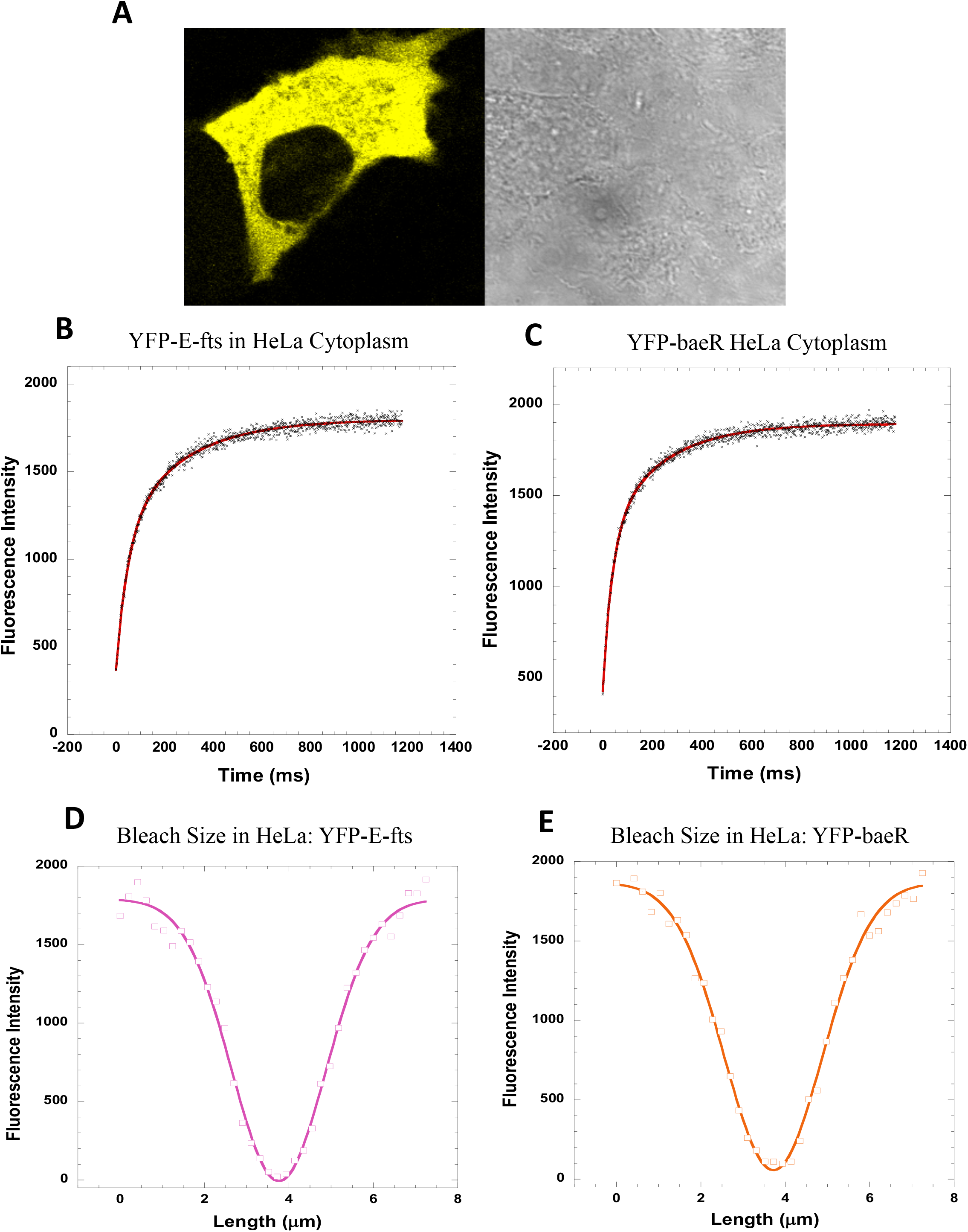
(A) A typical HeLa cell after successful micro-injection with a fluorescent labeled E-fts protein. (B-C): FRAP recovery profiles of (B) E-fts fused with YFP (N=43; R=0.99) (C) baeR fused with YFP (N=42; R=0.99) in HeLa cell cytoplasm. (D-E): Gaussian fits of averaged post-bleach size profiles for (D) E-fts fused with YFP (N=29; R=0.99) (E) baeR fused with YFP (N=36; R=0.99) in HeLa cell cytoplasm.

*E.coli* bacterial cells are too small for micro-injection of proteins inside the cell(Mika & Poolman, 2011). Hence, baeR-YFP and E-fts-YFP proteins were expressed in the bacterial cell cytoplasm. To avoid fluorescence signal saturations, we have calibrated protein expression levels using the *E.coli tuner* cells(Hartinger et al., 2010), where protein expression is tightly regulated. A typical example of *E.coli* cells expressing YFP fused E-fts protein is shown in Figure 7A. In Figure 7B and C FRAP recovery profiles of E-fts fused with YFP and baeR fused with YFP in *E.coli* cytoplasm are shown. Figures 7D and E show the post-bleach fluorescens intensity profiles, from which r_e_ values were calculated (Table S2). For the E-fts fused to YFP *D* = 1.6 μm^2^s^−1^ and for baeR fused to YFP *D* =0.95 μm^2^s^−1^ were calculated (Table S4). In addition, we have measured the diffusion coefficient of the YFP protein by itself in the cytoplasm of *E.coli tuner* cells, resulting in a value of *D* = 4.6 μm^2^s^−1^. Previously, a value of 7.7±2.5 μm^2^s^−1^ was determined for GFP in *E.coli* D5Hα cells using photobleaching and photo activation using conventional FRAP(Elowitz et al., 1999). It was further reported that GFP diffusion rates depend on the expression level, with a value of *D*~ 3.6±0.7 μm^2^s^−1^ being reported at higher protein expression. Furthermore, insertion of a histidine tag during construct designing resulted in *D*~ 4.0±2.0 μm^2^s^−1^ (Elowitz et al., 1999). The published values indicate that the Line-FRAP method predicts correctly diffusion coefficients in prokaryotic cells. Moreover, the data qualities generated using Line-FRAP is less noisy than those shown in previous reported FRAP measurements in *E.coli*(Elowitz et al., 1999; Kumar et al., 2010).

**Figure 7:**
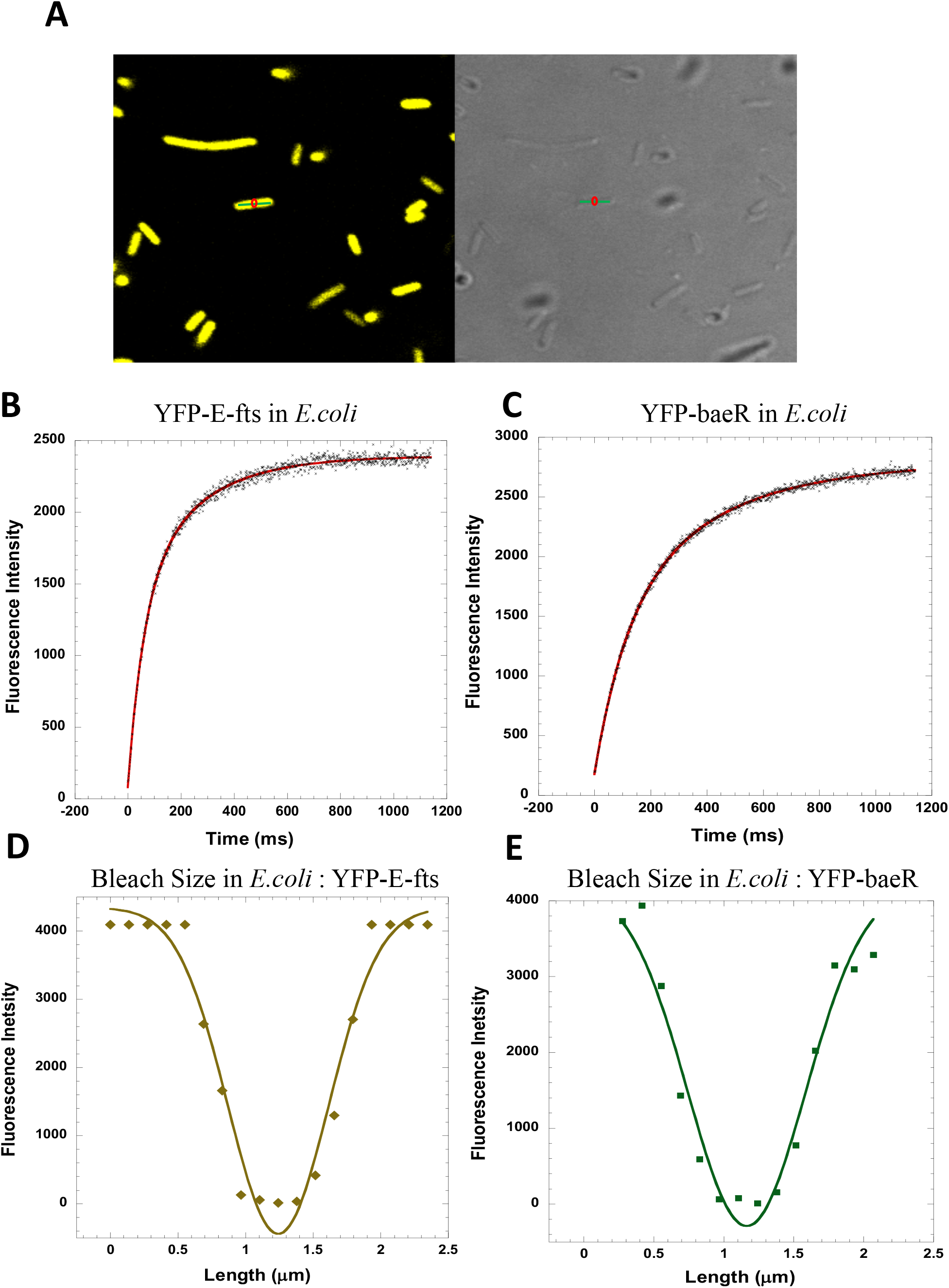
(A) Typical E.Coli cells after expression of YFP fused E-fts protein. (B, C): FRAP recovery profiles of (B) E-fts fused with YFP (N=49; R=0.99) (C) baeR fused with YFP (N=52; R=0.99) in E.coli cytoplasm. (D, E): Gaussian fits of post bleach size profiles for (D) E-fts fused with YFP (R= 0.98) (E) baeR fused with YFP (R= 0.97) in E.coli cytoplasm.

Figure 8 provides a comparison of diffusion coefficient values for E-fts and baeR in different environments as determined using Line-FRAP. Figure S4 shows a comparison of (A) τ_1/2_ of all corresponding diffusion processes and (B) percentage of recovery after FRAP. Fusing YFP doubles their MW, and subsequently the diffusion coefficients were reduced in PBS buffer solution. Both the proteins with YFP fusions showed further reduced mobility in the complex environment of the HeLa cell cytoplasm. It is known that mammalian cell cytoplasm is 2-3 times as dense as aqueous buffer solution(Dix & Verkman, 2008; Mika & Poolman, 2011). Prokaryotic cell cytoplasm are further packed, with diffusion rates being over 10-fold slower than in buffer(Mika & Poolman, 2011). The fold-change in diffusion coefficients from PBS to HeLa to *E.coli* is also in line with the expected reduction due to the increased viscosity(Dix & Verkman, 2008; Mika & Poolman, 2011).

**Figure 8:**
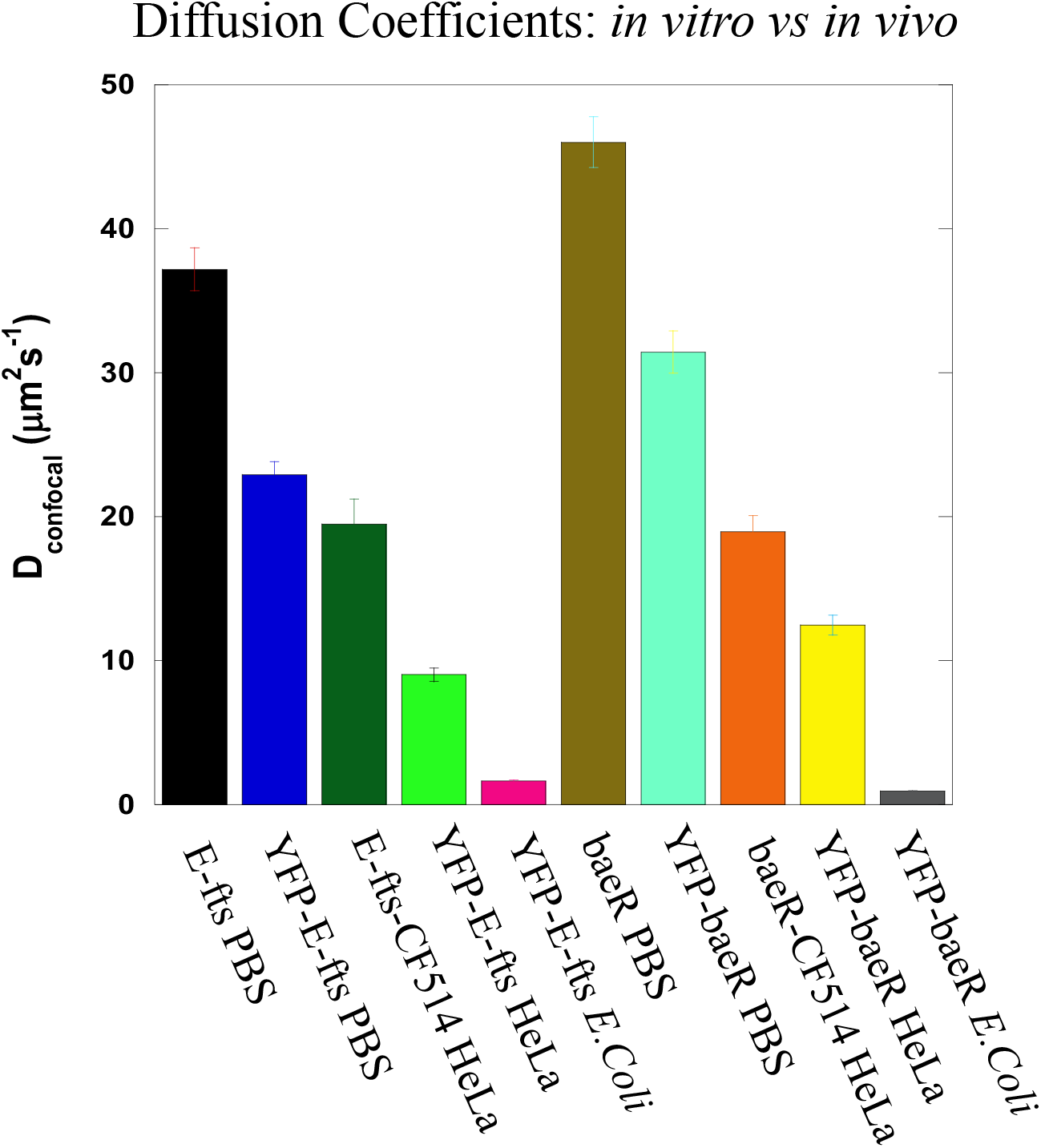
Comparing diffusion coefficients of E-fts and baeR proteins chemically labeled or fused with YFP in PBS buffer, HeLa cell cytoplasm and E.coli bacteria cells. (Final error bars are calculated after combining all the individual errors from the respective average fits).

To further test the validity of our Line-FRAP method, we have carried out Diffusion rate measurements in *E.coli* cells under osmotically challenging environments. It is well known that at higher osmotic stress, the cell wall starts to stretch causing the cytoplasm to shrink(Konopka et al., 2009; Mika et al., 2014). As a consequence of the increased crowding density inside the bacterial cell cytoplasm, the mobility of proteins slows down, which is reflected in lower diffusion coefficient values(Konopka et al., 2009; Mika et al., 2014; Sochacki et al., 2011). Figure 9A and B show normalized FRAP recovery profiles and corresponding post-bleach profiles for YFP in *E.coli* under different osmotic stress conditions. Figure 9C shows a comparison of the diffusion coefficient values under different osmotic conditions. Figure S4C shows a comparison of τ_1/2_ of all corresponding diffusion processes and Figure S4D shows the percentage of recovery after FRAP. Increasing osmolarity was done by replacing the 2TY medium to HEPES buffer plus varying concentrations of NaCl. This resulted in decreasing *D*_confocal_ values, from *D* = 4.6 μm^2^s^−1^ in 2TY medium (including 85 mM NaCl) to 3.2, 2.4 and 1.9 μm^2^s^−1^ in presence of 150, 300 and 500 mM NaCl respectively (Table S5). Detailed visual images of *E.Coli* cells under osmotically challenging environments are provided in the supporting information Figure S5 (A-D). At very high concentration of salt (1000 mM NaCl), almost no recovery of YFP protein was observed inside the *E.coli* cells (Figure S5D). It suggest that addition of excess salt concentration may leads to protein aggregation or formation of diffusive barriers leading to compartmentalization of YFP, which contributed towards the immobility of YFP in *E.Coli*. (Van Den Bogaart et al., 2007). These results confirm the power of the here presented method in challenging conditions.

**Figure 9:**
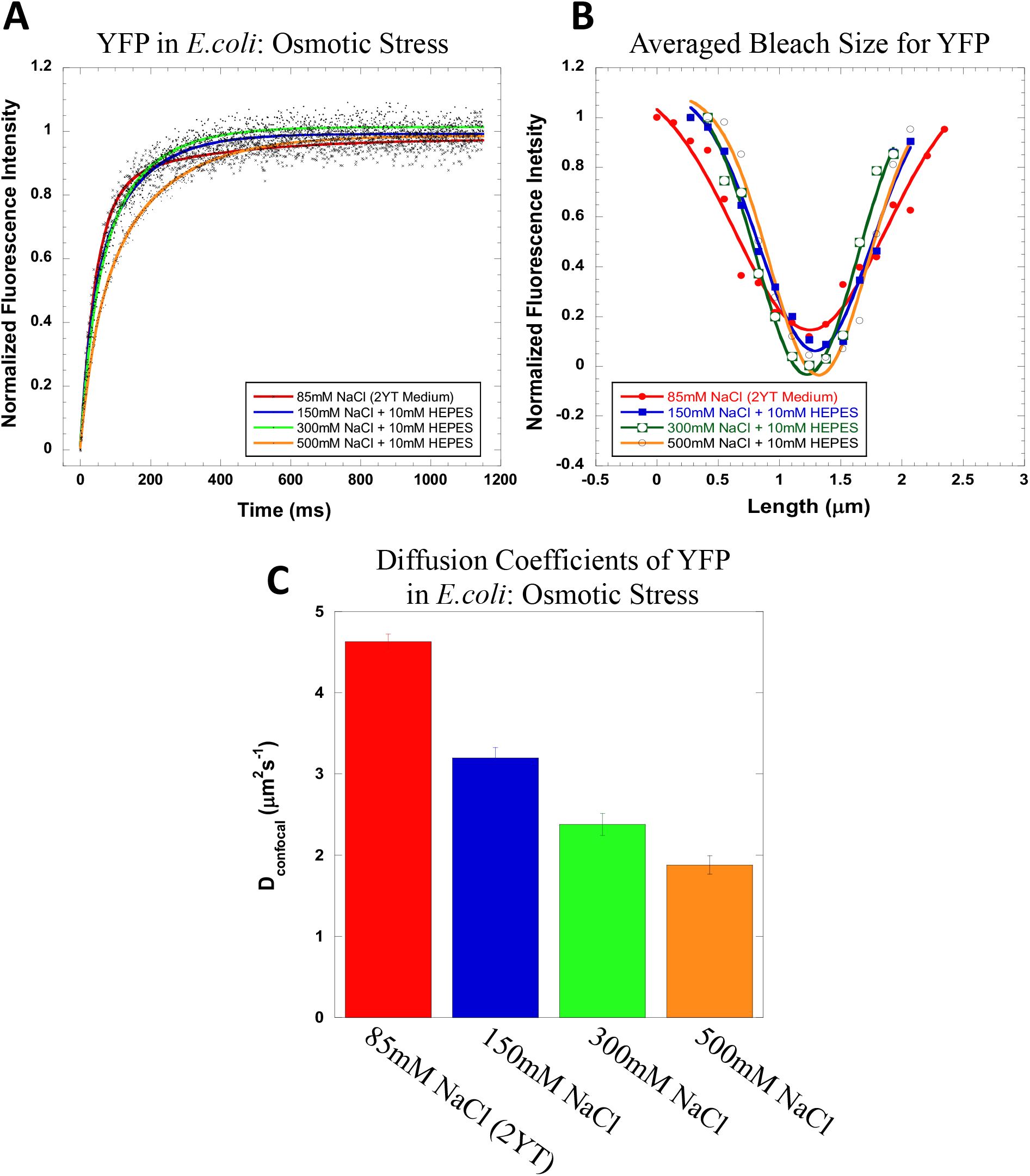
(A) Normalized FRAP recovery profiles (N=20; R=0.97 for YFP in 2YT, N=30; R=0.98 in 150mM, N=30; R=0.99 in 300mM and N=30; R=0.99 in 500mM NaCl). (B) Corresponding Gaussian fits of averaged post-bleach size profiles (R=0.98 for YFP in 2YT, R=0.99 for 150mM, 300mM and 500mM NaCl). And (C) Comparison of diffusion coefficients for YFP protein expressed in E.coli cells under different osmotic stress environments (final error bars are calculated after combining all the individual errors from the respective average fits).

Our data show that we have developed a universal method by which one can compare diffusion coefficients measurements in different environments including *in vivo* by using FRAP. This allows diffusion measurements to be done using a simple, commonly available setup and applying straight forward data analysis tools. Moreover, protein-concentrations used for these measurements are in the μM range, which is in line with *in vivo* protein concentrations. The main advantage of the here presented method is the much faster data acquisition rate, which provides exact measurements of the bleach area already 1 ms after bleach, in comparison to 20-100 ms in conventional FRAP. For fast diffusion molecules, including proteins, this time difference is crucial to obtain exact values of the bleach diameter (which is squared in equation 3) and recovery half-times, which together provide reliable diffusion coefficients. We show that using the same proteins but conventional FRAP, the measurements of these parameters is vastly different (due to the slow frame rate), which results in inconsistent diffusion rates for fast moving molecules. Overall, the here presented system has major advantages over other methods, both in usability and versatility.

## Experimental

### Materials and Methods

Dulbecco’s phosphate buffered saline (PBS 1X) was purchased from Biological Industries (catalog no. 02-023-1A). CF514 fluorescent labelling dye was purchased from Biotium. All other reagents used are described in details in a table format (Table 2, Key resources)

**Table 2:**
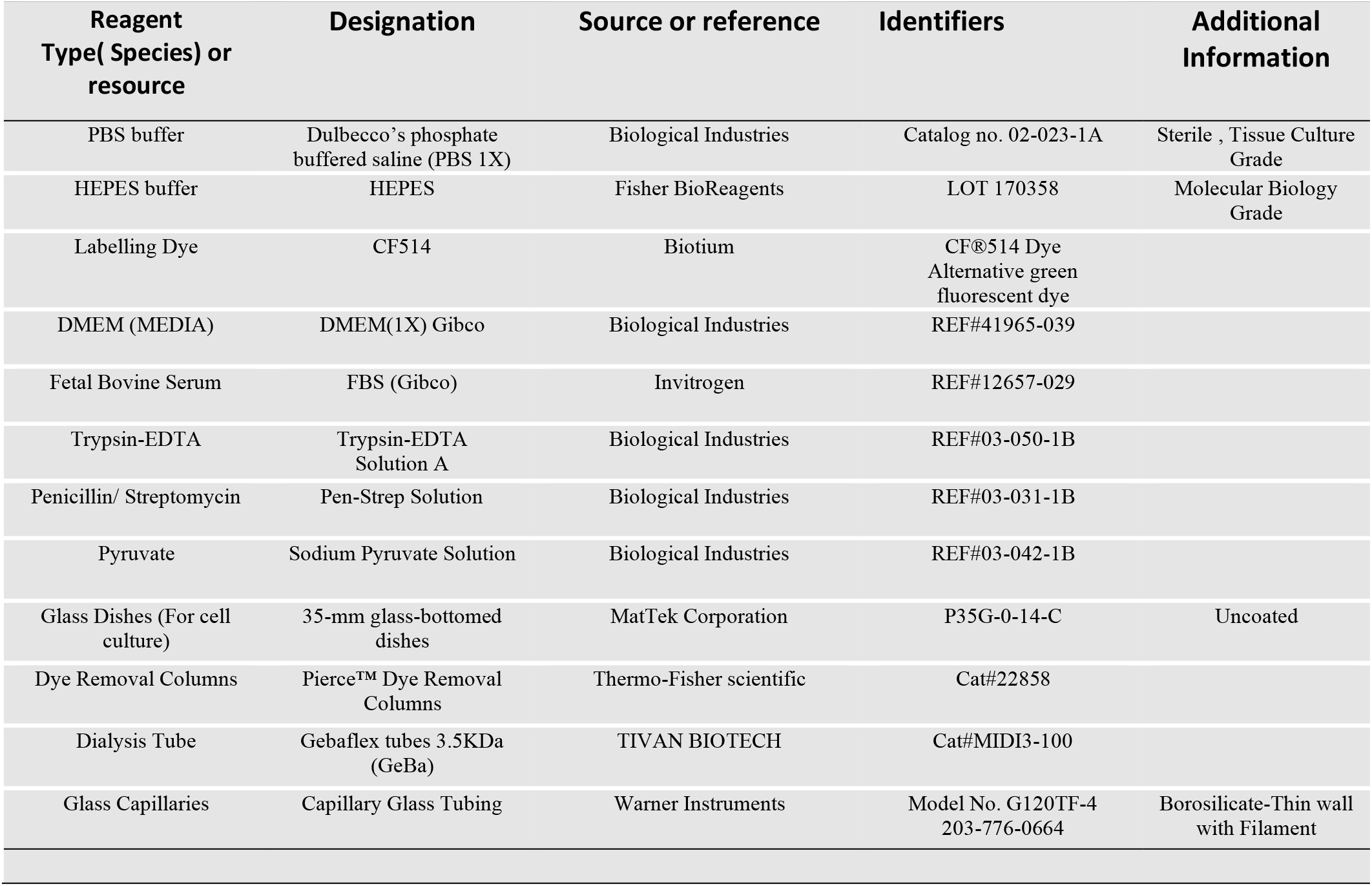
Key resources

#### Mammalian Cell culture

HeLa cells were grown in 35-mm glass-bottomed dishes (MatTek Corporation) in DMEM (Invitrogen) supplemented with 1x pyruvate, penicillin/streptomycin (BioLabs), and 10% fetal bovine serum (Equitech-BIO. Inc.). The cells were subcultured when 80% confluence was reached using trypsin-EDTA for cell detachment. 2×10^5^ HeLa cells in 2.5 ml of DMEM were pipetted into glass bottomed dishes and incubated overnight. The cells were cultured in humid atmosphere at 37 °C and 5% CO_2_. The cells were imaged 24–30 h after the seeding process. Prior to the microinjection, the medium was aspirated, and the fresh medium was supplemented with 25 mM HEPES, pH 7.4.

#### Cloning and protein expression in E.coli

Three different constructs were cloned for BaeR and Efts. BaeR or Efts fused in their N-terminus to YFP (with A206K mutation that prevents self-dimerization) with a linker were cloned to pET21a plasmid. The same construct was cloned to pET28-his-sumo plasmid for purification. In addition, BaeR or Efts were cloned to pET28-his-sumo plasmid without YFP fusion. Cloning was done by RF cloning and by using a synthesized gene of YFP A206K (Twist biosciences). The YFP A206K BaeR or Efts in pET21a was transformed to Tuner™(DE3) Competent Cells (Novagene) which allow adjustable levels of protein expression throughout all cells in a culture. They were grown in 37°C until they reached OD600= 0.6 and induced by adding IPTG to a final concentration of 0.1 mM in 37°C for three hours-then stored at 4°C until imaging and measuring diffusion. For higher osmotic stress conditions, the YFP expressed *E.coli* bacteria cells were centrifuged, and resuspended in 10 mM HEPES buffer solution (pH 7.2) with different NaCl salt concentrations (150-500 mM). The incubation time in buffer solutions after resuspension was varied between 30 mins to 3 hours, which had to effect on the calculated diffusing rates. For protein-purification, YFP A206K BaeR or Efts in pET28-his-sumo and BaeR or Efts in pET28-his-sumo plasmid were transformed to BL21 (DE3) competent cells. They were grown at 37°C until they reached OD_600_= 0.6-0.8 before adding 0.1 mM IPTG for protein production in 16°C overnight.

#### Protein Purification

After overnight growth, the cells were harvested for 10 minutes at 5000 rpm and stored at −20°C. The pellets were resuspended in 50 mM Tris-HCl, 200 mM NaCl and 10 mM Imidazole pH=8 buffer. Protease inhibitor, lysozyme and benzonase were added. After homogenization, sonication was done by using 3 cycles of 10 seconds on and 30 seconds off keeping the sample on ice. Then, the samples were centrifuged at 14000 rpm for 60 minutes at 4°C. The supernatant was used for further purification. Ni-NTA gravity columns of 200 ul Ni-NTA (Merck) were equilibrated with binding buffer containing 50 mM Tris-HCl pH=8, 200 mM NaCl, 10 mM Imidazole and 10% glycerol. The supernatant was loaded onto the column after 0.45 nm filtration (Millex-HV; Millipore), then, washed with 50 mM Tris-HCl, 200 mM NaCl, 20 mM Imidazole and 10% glycerol twice. On column cleavage was done by sumo-protease in elution buffer: 50 mM Tris-HCl, 200 mM NaCl, 10% glycerol, 1mM DTT, 1:200 sumo protease. The cleavage was done one hour in room temp and then overnight at 4°C under constant rotation. Elution from beads was done twice using the cleavage buffer. For BaeR and Efts this was followed by anion exchange of a FF Q column (GE). Purified proteins were analyzed by SDS-PAGE gel-electrophoresis and its concentrations were determined using a nanodrop. The purified protein was dialyzed using midi-gebaflex 3.5 kDa (Geba) against 0.1M NaHCO3 buffer that is compatible to labelling with CF514 dye. The protein was then aliquoted, flash-frozen in liquid nitrogen, and stored at −80°C.

#### Protein labelling with CF514

The purified BaeR and Efts were labelled with CF514 dye (Biotium) according to the manufacturer recommendations. Labelled proteins were dialyzed against PBS using Gebaflex 3.5KDa (GeBa) and further purified from the dye using Pierce™ Dye Removal Columns (Cat#22858, Thermo-Fisher scientific).

#### Microinjection into HeLa cells

Microinjections were performed using Eppendorf FemtoJet microinjector attached to Eppendorf InjectMan NI2 micromanipulator. The different fluorescent proteins were dissolved in PBS and injected into cells using glass capillaries from Warner instruments and pulled by vertical puller (Narishige). For every measurement, a single pressure pulse was applied to deliver the sample into the cell. Air was administrated at 15–25 hPa for 0.1-0.3 s. For injections, single cells containing morphologically healthy and well-connected HeLa cells were selected. Before and after the microinjection, cell morphology and the membrane integrity were confirmed by visually inspecting the injected cells.

#### Confocal microscopy and FRAP Analysis

##### Confocal microscopy

Imaged were collected with an Olympus IX81 FluoView FV1000 Spectral/SIM Scanner confocal laser-scanning microscope, using 60X DIC oil-immersion objective, N.A. 1.35. For CF514/YFP fluorescence measurements, excitation was done at 515-nm, using a diode laser at an output power of 3% and 1% for low and high YFP/CF514 dye concentrations. Emission was recorded from 530 to 630 nm using the spectral detection system. All image analyses were performed using FluoView software and data analyses were performed using Kaleidagraph software version 4.1 (Synergy).

#### Line-FRAP

Line-FRAP was carried out in liquid drops, HeLa cells or *E.coli.* The target cells were imaged with the main scanner at 515-nm excitation using 3% of the maximal intensity applying a 60x differential interference contrast oil-immersion objective lens. For photobleaching, a “tornado” of 4 pixels diameter was used in the simultaneous stimulus scanner. The circle area of the bleach was kept exactly in the middle of the scanning line. The unidirectional lines were scanned with time intervals of 1.256 ms 1000 times (equivalent to 1.256 s). The number of scans before, during, and after photobleaching was 10, 42, and 948, respectively. Photobleaching was achieved with the simultaneous laser at 405-nm excitation for 10 or 63 milisecond duration, using full intensity. The simultaneous scanner moved at a speed of 100 μs/pixel to perform an efficient photobleach. We have used two simultaneous scanners during the FRAP experiments: one scanner (at 405 nm) for photobleaching and another scanner (at 515 nm) for data acquisition. We have varied the intensity of the lasers to achieve good signal/noise ratio. Fluorescence recovery plots were fitted to a double exponent growth curve. To do a comparative study, FRAP experiments were also performed inside the PBS buffer drops putting a cover slip upside like in a sandwich mode.

## ACKNOWLEDGMENT

We would like to thanks Dr. Hagen Hofmann from the department of structural biology, Weizmann Institute of Science for advice and critically reading the manuscript, and Dr. Vladimir Kiss from the department of biomolecular sciences for his help in the microscopy work. The work was supported by a grant of the Israel Science Foundation number 1268/18 and a grant by the United States - Israel Binational Science Foundation number 2015376.

## Supporting Information

**Table S1:** Comparison of Line-FRAP with other Fluorescence microscopy techniques for measuring diffusion rates of Proteins in buffer and living cells

| Techniques | Principle | Necessary Requirements | Sensitivity | Pros/Cons |
| --- | --- | --- | --- | --- |
| **FCS** | Auto-correlation of fluorescence fluctuations | Low fluorophore concentration with high quantum yield | Not sensitive to mild and uniform bleaching | Works mainly in ideal solution condition (Gold standard) but blind towards immobile fraction if any |
| **RICS** | combination of the temporal scales of single point FCS with the spatial information obtained from ICS | Similar to FCS | Pixel size is an important consideration in designing an experiment | Indirectly measure binding by detecting changes in the diffusion coefficients *in vivo* (multiple phenomenon detections possible within a cell). Limited towards the processes that are very slow (i.e., seconds timescale) and thus cannot show correlation between pixels of one frame |
| **SPT** | Localization of a fluorescent molecule is traced over time | Low fluorophore concentrations | Sensitive to photo-bleaching and blinking | Discriminate between normal and anomalous diffusion. Auto fluorescence from the background can interfere with the measuring process. |
| **FSM** | Trajectories of co-localized fluorophore cluster inside macromolecular structures | Stable speckle formation is necessary. Primarily used to investigate the dynamics of polymers. | Less sensitive to photo-bleaching and blinking than SPT | Similar to SPT but not suitable for studying rapidly moving monomeric proteins as they do not form stable speckles. |
| **PIPE** | Spreading of initially localized photo-converted fluorophores | Photo-convertible fluorophores | Not sensitive to mild and uniform bleaching | can measure rapidly moving proteins in cytoplasm even at high concentrations |
| **FRAP** | Recovery of averaged fluorescence signal in photo-bleached region | Calibration of beam width, High concentration of fluorophores | Sensitive to reversible Photo-bleaching, bleach pulse operating time scale | Easy to perform and analysis in conventional LSMs but Limited towards the fast moving molecules especially *in vitro* condition. |
| **FDAP/FLAC** | Decay of average fluorescence signal after photo activation by a focused laser flash | Photo-convertible fluorophores, calibration of beam width same as FRAP | Sensitive to photo-bleaching | FDAP can be complementary to FRAP study |
| **LINE-FRAP** | Recovery of averaged fluorescence through a single line (along a pixel) in photo-bleached region | Two simultaneous scanners, very short bleaching point/area | Sensitive to reversible photo-bleaching, bleach time scale, bleach size area | Capable of measuring diffusion rates in solution, Eukaryotic, and Prokaryotic |

RICS= Raster image correlation spectroscopy; FSM= fluorescence speckle microscopy; FLAC= Fluorescence loss after photo-conversion; FDAP=Fluorescence decay after photo-conversion;

**Table S2:** Diffusion Coefficients calculated from classical XY-FRAP data for labelled BSA in PBS buffer at different bleach time durations with variations in frame acquisition speed

| Frame Rate | Bleach Pulse durations | $k_1$<br>$\text{ms}^{-1}$ | $k_2$<br>$\text{ms}^{-1}$ | $a_1$ | $a_2$ | $p_1$ | $p_2$ | $\tau_1$ | $\tau_2$ | $\tau_{1/2}$<br>(ave) | $c_n$<br>$\mu\text{m}$ | $r_n$<br>$\mu\text{m}$ | $c_e$<br>$\mu\text{m}$ | $r_e$<br>$\mu\text{m}$ | $D_{\text{confocal}}$<br>$\mu\text{m}^2 \text{s}^{-1}$ |
| --- | --- | --- | --- | --- | --- | --- | --- | --- | --- | --- | --- | --- | --- | --- | --- |
| 72 X 72 Pixel<br>( 14.90 $\mu\text{m}$<br>X 14.90 $\mu\text{m}$) | 63 ms | 0.011 | 0.001 | 212.5 | 104 | 0.67 | 0.33 | 0.063 | 0.693 | 0.0898 | 0.92 | 1.27 | 4.42 | 3.20 | 29.4 |
| 36 X 36 Pixel<br>( 7.45 $\mu\text{m}$<br>X 7.45 $\mu\text{m}$) | 63 ms | 0.031 | 0.003 | 612.5 | 169 | 0.78 | 0.22 | 0.022 | 0.231 | 0.0278 | 0.92 | 1.27 | 4.28 | 3.10 | 89.6 |
| 36 X 36 Pixel<br>( 7.45 $\mu\text{m}$<br>X 7.45 $\mu\text{m}$) | 10 ms | 0.039 | $1 \times 10^{-5}$ | 799 | 790 | 0.50 | 0.50 | 0.018 | 69.3 | 0.035 | 0.54 | 0.74 | 2.35 | 1.70 | 21.4 |

The rate constants ( $k$ ) and amplitudes ( $a$ ) are from the double-exponential fit of the FRAP data. The $c$ values were obtained from Gaussian fits using equation 5. The respective radius ( $r$ ) are calculated assuming the respective bleach circles diameter lies at 60% height from the bottom of the Gaussian Profiles ( $r=1.38*c$ ).

**Table S3:**
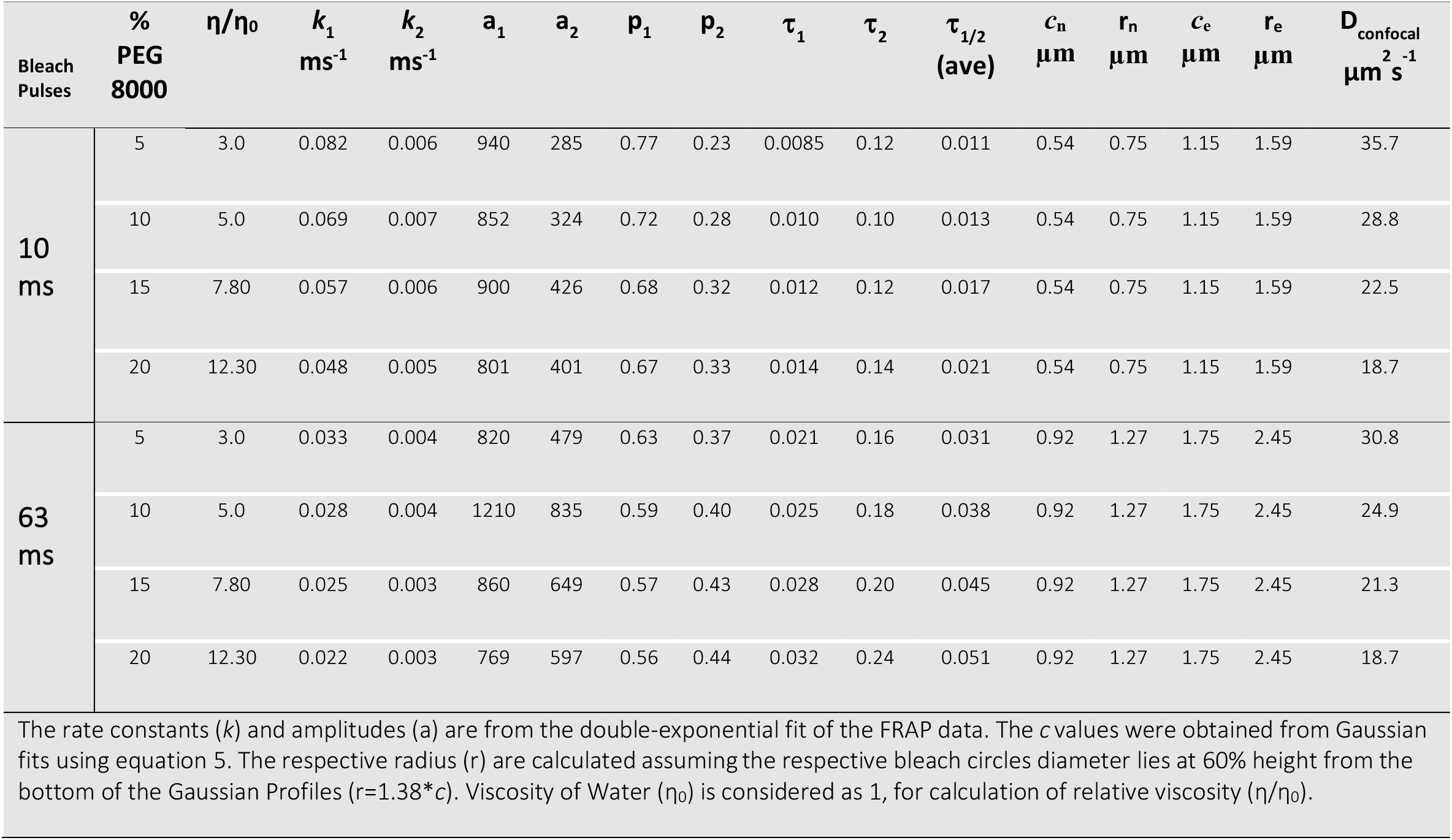
Diffusion Coefficients calculated from FRAP data for labelled BSA in PBS buffer in the presence of PEG8000 as Crowder

| Bleach Pulses | % PEG 8000 | $\eta/\eta_0$ | $k_1$<br>$\text{ms}^{-1}$ | $k_2$<br>$\text{ms}^{-1}$ | $a_1$ | $a_2$ | $p_1$ | $p_2$ | $\tau_1$ | $\tau_2$ | $\tau_{1/2}$<br>(ave) | $c_n$<br>$\mu\text{m}$ | $r_n$<br>$\mu\text{m}$ | $c_e$<br>$\mu\text{m}$ | $r_e$<br>$\mu\text{m}$ | $D_{\text{confocal}}$<br>$\mu\text{m}^2 \text{s}^{-1}$ |
| --- | --- | --- | --- | --- | --- | --- | --- | --- | --- | --- | --- | --- | --- | --- | --- | --- |
| 10 ms | 5 | 3.0 | 0.082 | 0.006 | 940 | 285 | 0.77 | 0.23 | 0.0085 | 0.12 | 0.011 | 0.54 | 0.75 | 1.15 | 1.59 | 35.7 |
|  | 10 | 5.0 | 0.069 | 0.007 | 852 | 324 | 0.72 | 0.28 | 0.010 | 0.10 | 0.013 | 0.54 | 0.75 | 1.15 | 1.59 | 28.8 |
|  | 15 | 7.80 | 0.057 | 0.006 | 900 | 426 | 0.68 | 0.32 | 0.012 | 0.12 | 0.017 | 0.54 | 0.75 | 1.15 | 1.59 | 22.5 |
|  | 20 | 12.30 | 0.048 | 0.005 | 801 | 401 | 0.67 | 0.33 | 0.014 | 0.14 | 0.021 | 0.54 | 0.75 | 1.15 | 1.59 | 18.7 |
| 63 ms | 5 | 3.0 | 0.033 | 0.004 | 820 | 479 | 0.63 | 0.37 | 0.021 | 0.16 | 0.031 | 0.92 | 1.27 | 1.75 | 2.45 | 30.8 |
|  | 10 | 5.0 | 0.028 | 0.004 | 1210 | 835 | 0.59 | 0.40 | 0.025 | 0.18 | 0.038 | 0.92 | 1.27 | 1.75 | 2.45 | 24.9 |
|  | 15 | 7.80 | 0.025 | 0.003 | 860 | 649 | 0.57 | 0.43 | 0.028 | 0.20 | 0.045 | 0.92 | 1.27 | 1.75 | 2.45 | 21.3 |
|  | 20 | 12.30 | 0.022 | 0.003 | 769 | 597 | 0.56 | 0.44 | 0.032 | 0.24 | 0.051 | 0.92 | 1.27 | 1.75 | 2.45 | 18.7 |

The rate constants ( $k$ ) and amplitudes ( $a$ ) are from the double-exponential fit of the FRAP data. The $c$ values were obtained from Gaussian fits using equation 5. The respective radius ( $r$ ) are calculated assuming the respective bleach circles diameter lies at 60% height from the bottom of the Gaussian Profiles ( $r=1.38*c$ ). Viscosity of Water ( $\eta_0$ ) is considered as 1, for calculation of relative viscosity ( $\eta/\eta_0$ ).

**Table S4:** Diffusion Coefficients calculated from FRAP data for different labelled proteins in different environments

| Proteins | Environ<br>ment | $k_1$<br>$ms^{-1}$ | $k_2$<br>$ms^{-1}$ | $a_1$ | $a_2$ | $p_1$ | $p_2$ | $\tau_1$ | $\tau_2$ | $\tau_{1/2}$<br>(ave) | $c_n$<br>$\mu m$ | $r_n$<br>$\mu m$ | $c_e$<br>$\mu m$ | $r_e$<br>$\mu m$ | $D_{confocal}$<br>$\mu m^2 s^{-1}$ |
| --- | --- | --- | --- | --- | --- | --- | --- | --- | --- | --- | --- | --- | --- | --- | --- |
| E-fts | PBS<br>labeled<br>with CF514<br>Dye | 0.041 | 0.005 | 887 | 482 | 0.65 | 0.35 | 0.017 | 0.14 | 0.024 | 0.92 | 1.27 | 1.73 | 2.38 | 37.2 |
|  | PBS<br>fused with<br>YFP | 0.032 | 0.005 | 780 | 493 | 0.61 | 0.39 | 0.022 | 0.14 | 0.032 | 0.92 | 1.27 | 1.50 | 2.07 | 22.9 |
|  | HeLa<br>Cytoplasm<br>labeled<br>with CF514<br>Dye | 0.030 | 0.004 | 880 | 611 | 0.59 | 0.41 | 0.023 | 0.17 | 0.036 | 0.92 | 1.27 | 1.45 | 2.00 | 19.5 |
|  | HeLa<br>Cytoplasm<br>fused with<br>YFP | 0.020 | 0.004 | 753 | 674 | 0.53 | 0.47 | 0.035 | 0.17 | 0.056 | 0.92 | 1.27 | 1.13 | 1.55 | 9.0 |
|  | *E.coli*<br>fused with<br>YFP | 0.017 | 0.004 | 1294 | 1018 | 0.56 | 0.44 | 0.042 | 0.17 | 0.063 | 0.54 | 0.75 | 0.38 | 0.52 | 1.6 |
| baeR | PBS<br>labeled<br>with CF514<br>Dye | 0.051 | 0.006 | 786 | 431 | 0.65 | 0.36 | 0.014 | 0.12 | 0.020 | 0.92 | 1.27 | 1.73 | 2.38 | 46.0 |
|  | PBS<br>fused with<br>YFP | 0.041 | 0.006 | 985 | 469 | 0.68 | 0.32 | 0.017 | 0.13 | 0.023 | 0.92 | 1.27 | 1.5 | 2.07 | 31.4 |
|  | HeLa<br>Cytoplasm<br>labeled<br>with CF514<br>Dye | 0.028 | 0.004 | 468 | 295 | 0.61 | 0.39 | 0.025 | 0.17 | 0.037 | 0.92 | 1.27 | 1.45 | 2.00 | 19.0 |
|  | HeLa<br>Cytoplasm<br>fused with<br>YFP | 0.026 | 0.005 | 835 | 632 | 0.57 | 0.43 | 0.027 | 0.15 | 0.041 | 0.92 | 1.27 | 1.15 | 1.59 | 12.5 |
|  | *E.coli*<br>fused with<br>YFP | 0.009 | 0.003 | 1246 | 1368 | 0.48 | 0.52 | 0.077 | 0.23 | 0.12 | 0.54 | 0.75 | 0.43 | 0.59 | 0.95 |
| YFP | *E.coli* | 0.023 | 0.004 | 1392 | 284 | 0.83 | 0.17 | 0.03 | 0.17 | 0.035 | 0.54 | 0.75 | 0.63 | 0.86 | 4.6 |

The rate constants ( $k$ ) and amplitudes ( $a$ ) are from the double-exponential fit of the FRAP data. The $c$ values were obtained from Gaussian fits using equation 5. The respective radius ( $r$ ) are calculated assuming the respective bleach circles diameter lies at 60% height from the bottom of the Gaussian Profiles ( $r=1.38*c$ ).

**Table S5:** Diffusion Coefficients of YFP protein in *E.coli* cytoplasm under Osmotic Stress Environment

| NaCl Conc.<br>(Environment) | $k_1$<br>$\text{ms}^{-1}$ | $k_2$<br>$\text{ms}^{-1}$ | $a_1$ | $a_2$ | $p_1$ | $p_2$ | $\tau_1$ | $\tau_2$ | $\tau_{1/2}$<br>(ave) | $c_n$<br>$\mu\text{m}$ | $r_n$<br>$\mu\text{m}$ | $c_e$<br>$\mu\text{m}$ | $r_e$<br>$\mu\text{m}$ | $D_{\text{confocal}}$<br>$\mu\text{m}^2 \text{s}^{-1}$ |
| --- | --- | --- | --- | --- | --- | --- | --- | --- | --- | --- | --- | --- | --- | --- |
| 85mM<br>(2YT Medium) | 0.023 | 1392 | 0.004 | 284 | 0.83 | 0.17 | 0.03 | 0.17 | 0.035 | 0.54 | 0.75 | 0.63 | 0.86 | 4.63 |
| 150mM<br>(10mM HEPES) | 0.033 | 642 | 0.008 | 853 | 0.43 | 0.57 | 0.02 | 0.087 | 0.037 | 0.54 | 0.75 | 0.46 | 0.62 | 3.20 |
| 300mM<br>(10mM HEPES) | 0.025 | 590 | 0.008 | 738 | 0.44 | 0.56 | 0.03 | 0.091 | 0.045 | 0.54 | 0.75 | 0.40 | 0.55 | 2.38 |
| 500mM<br>(10mM HEPES) | 0.023 | 590 | 0.006 | 972 | 0.38 | 0.62 | 0.03 | 0.126 | 0.057 | 0.54 | 0.75 | 0.40 | 0.55 | 1.87 |

The rate constants ( $k$ ) and amplitudes ( $a$ ) are from the double-exponential fit of the FRAP data. The $c$ values were obtained from Gaussian fits using equation 5. The respective radius ( $r$ ) are calculated assuming the respective bleach circles diameter lies at 60% height from the bottom of the Gaussian Profiles ( $r=1.38*c$ ).

**Figure S1:**
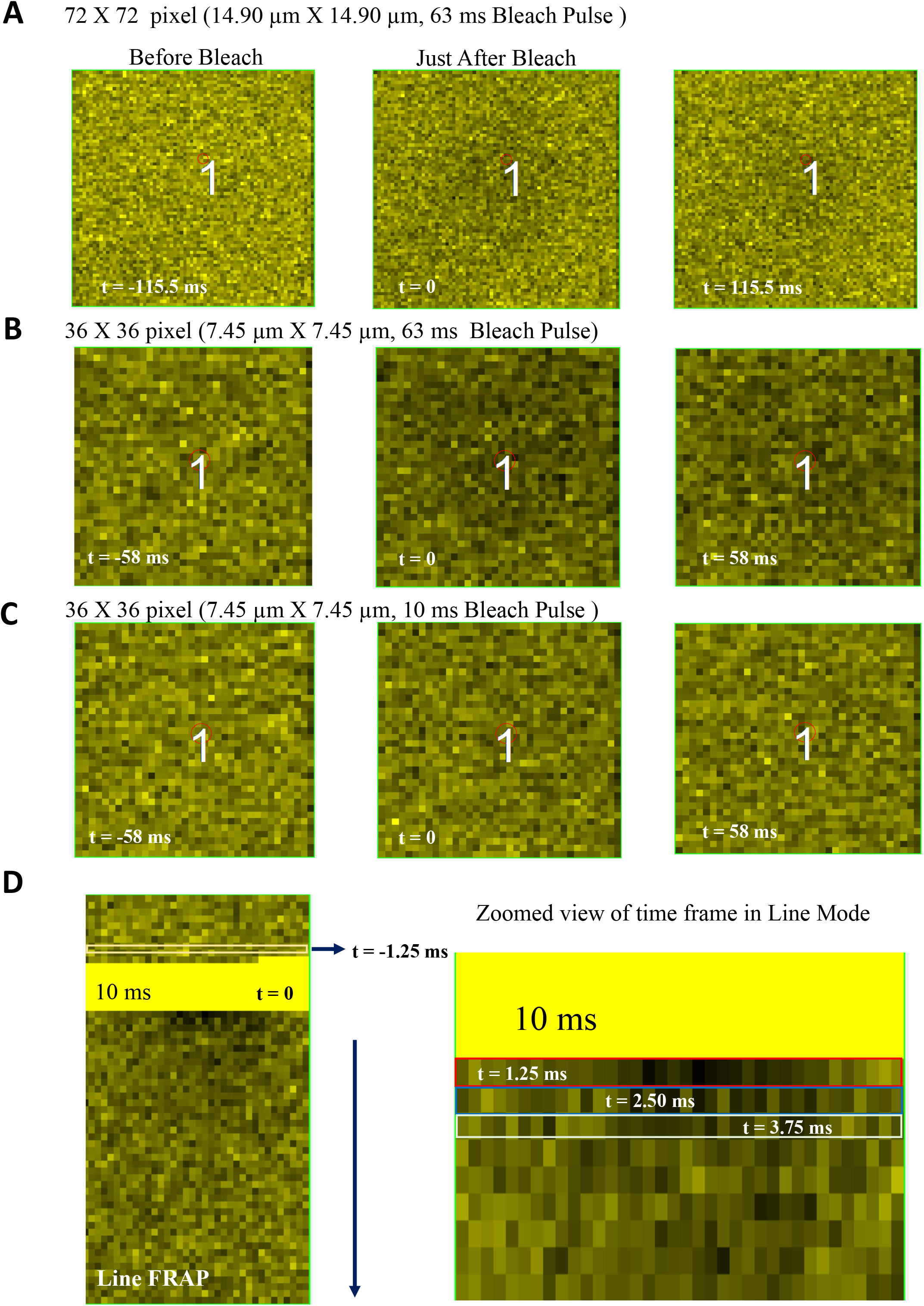
Acquisition of different Frame rates on fast diffusive molecule (BSA labeled with CF514 Dye in PBS buffer solution). Classical XY FRAP (A) 72X72 pixel (14.90 μm X 14.90 μm with 63 ms bleach pulse); (B) 36X36 pixel size (7.45 μm X 7.45 μm with 63 ms bleach pulse) and (C) with 10 ms bleach pulse; (D) Line FRAP with 1.25 ms time frame.

**Figure S2:**
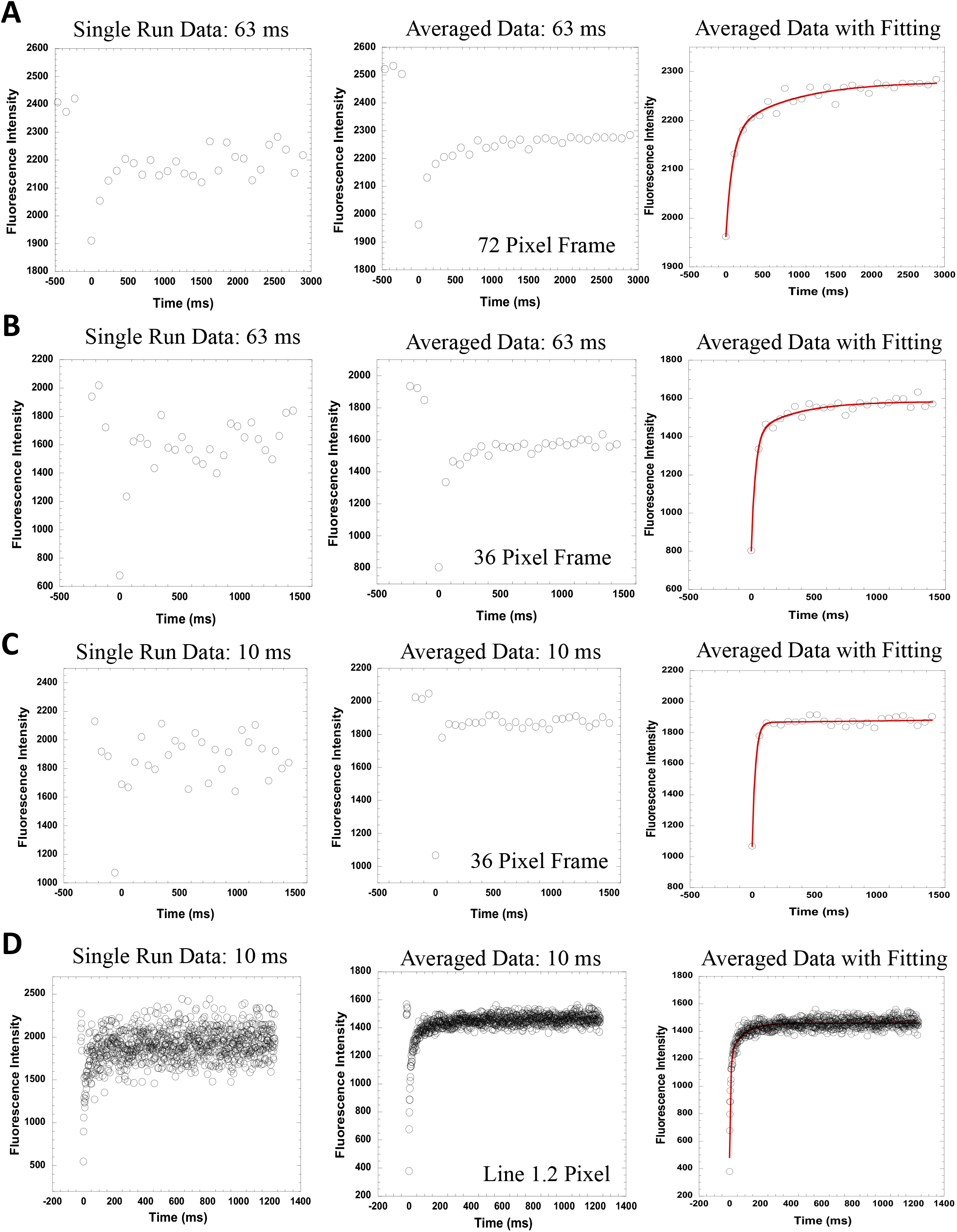
Effect of different frame rates on acquired recovery rate data quality in case of a faster diffusive molecule (BSA labeled with CF514 Dye in PBS buffer solution). Classical XY FRAP (A) 72X72 pixel (14.90 μm X 14.90 μm with 63 ms bleach pulse); (B) 36X36 pixel size (7.45 μm X 7.45 μm with 63 ms bleach pulse) and (C) with 10 ms bleach pulse; (D) Line FRAP with 1.25 ms time frame. Comparison of single run data vs averaged data (N=20) with their exponential fits are also shown in each cases.

**Figure S3:**
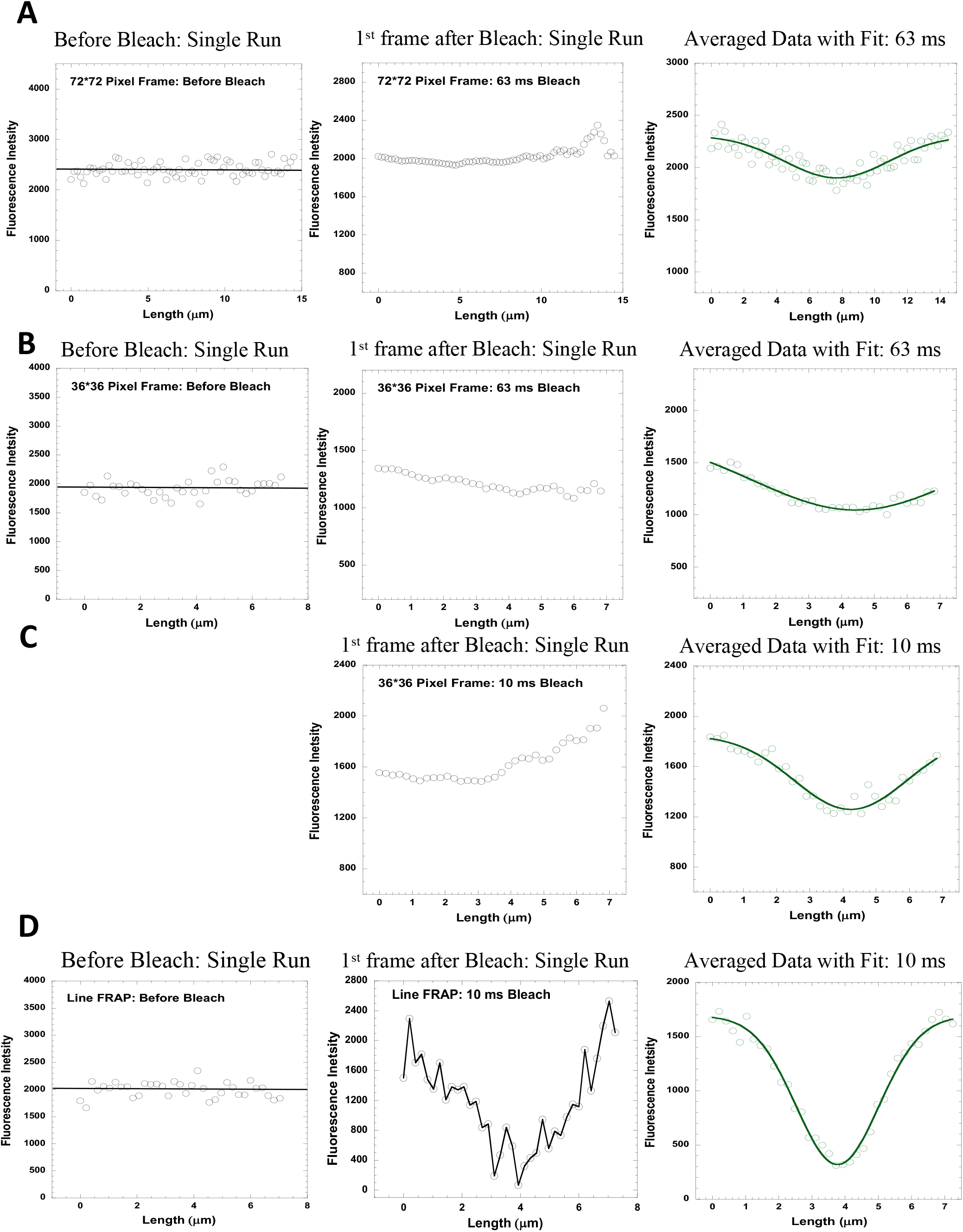
Effect of different frame rates on acquired bleach size data quality in case of a faster diffusive molecule (BSA labeled with CF514 Dye in PBS buffer solution). Classical XY FRAP (A) 72X72 pixel (14.90 μm X 14.90 μm with 63 ms bleach pulse); (B) 36X36 pixel size (7.45 μm X 7.45 μm with 63 ms bleach pulse) and (C) with 10 ms bleach pulse; (D) Line FRAP with 1.25 ms time frame. Comparison of single run data vs averaged data with Gaussian fitting (N=20) are also shown in each cases.

**Figure S4:**
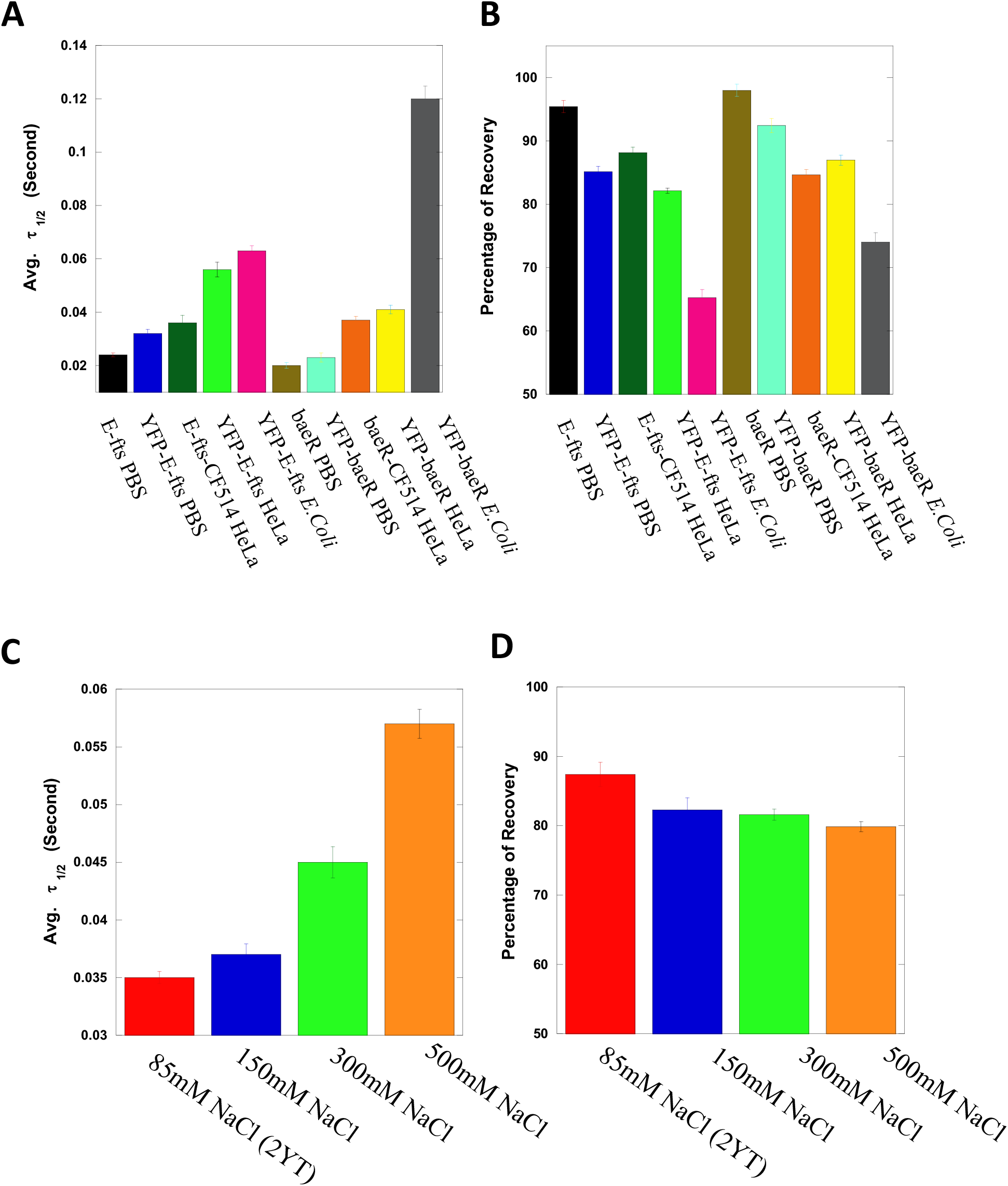
Comparison of (A) half-time (B) percentage of recoveries of E-fts and baeR proteins chemically labeled or fused with YFP in PBS buffer, HeLa cell cytoplasm or E.coli bacteria cells. (C) half-times (D) percentage of recoveries for YFP protein expressed in *E.coli* cells under different osmotic stress conditions.

**Figure S5:**
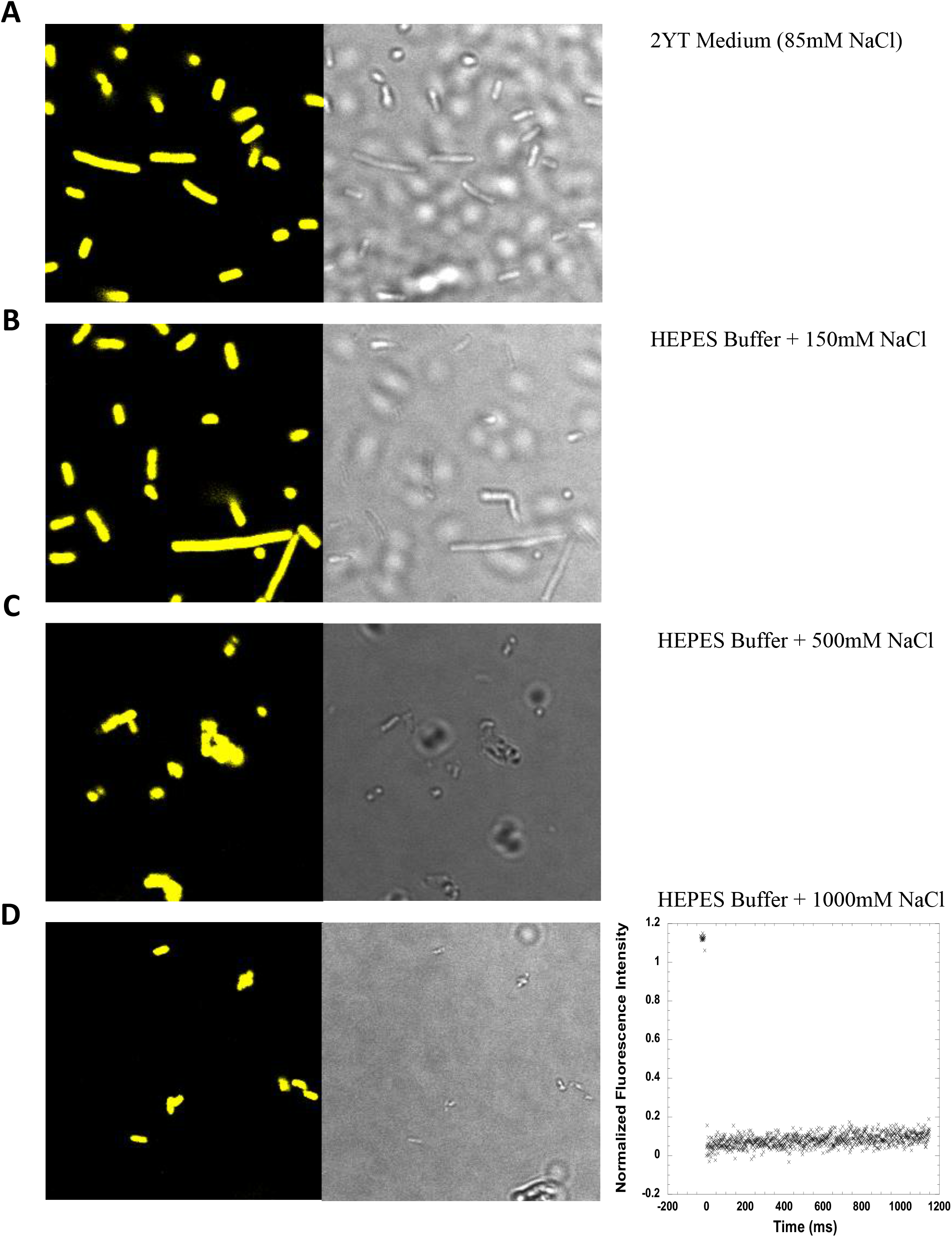
Visuals of *E.coli* cells under osmotically challenging environments. (A) 2YT medium (85 mM NaCl), (B) HEPES buffer + 150 mM NaCl, (C) HEPES buffer + 500 mM NaCl, (D) HEPES buffer + 1000 mM NaCl, where no fluorescence recovery was observed.

## REFERENCES

1. Goodsell, D. S. (1991). Inside a living cell. Trends in Biochemical Sciences, 16(C), 203–206. https://doi.org/10.1016/0968-0004(91)90083-8

2. Verkman, A. S. (2002). Solute and macromolecule diffusion in cellular aqueous compartments. In Trends in Biochemical Sciences (Vol. 27, Issue 1, pp. 27–33). https://doi.org/10.1016/S0968-0004(01)02003-5

3. Dix, J. A., & Verkman, A. S. (2008). Crowding Effects on Diffusion in Solutions and Cells. Annual Review of Biophysics, 37(1), 247–263. https://doi.org/10.1146/annurev.biophys.37.032807.125824

4. Agutter, P. S., Malone, P. C., & Wheatley, D. N. (2000). Diffusion theory in biology: A relic of mechanistic materialism. Journal of the History of Biology, 33(1), 71–111. https://doi.org/10.1023/A:1004745516972

5. Haugh, J. M. (2009). Analysis of reaction-diffusion systems with anomalous subdiffusion. Biophysical Journal, 97(2), 435–442. https://doi.org/10.1016/j.bpj.2009.05.014

6. Saxton, M. J. (2012). Wanted: A positive control for anomalous subdiffusion. In Biophysical Journal (Vol. 103, Issue 12, pp. 2411–2422). https://doi.org/10.1016/j.bpj.2012.10.038

7. Baumann, G., F. Place, R., & Foldes-Papp, Z. (2010). Meaningful Interpretation of Subdiffusive Measurements in Living Cells (Crowded Environment) by Fluorescence Fluctuation Microscopy. Current Pharmaceutical Biotechnology, 11(5), 527–543. https://doi.org/10.2174/138920110791591454

8. Lenzi, E. K., Ribeiro, H. V., Tateishi, A. A., Zola, R. S., & Evangelista, L. R. (2016). Anomalous diffusion and transport in heterogeneous systems separated by a membrane. Proceedings of the Royal Society A: Mathematical, Physical and Engineering Sciences, 472(2195). https://doi.org/10.1098/rspa.2016.0502

9. Schavemaker, P. E., Boersma, A. J., & Poolman, B. (2018). How important is protein diffusion in prokaryotes? In Frontiers in Molecular Biosciences (Vol. 5, Issue NOV). https://doi.org/10.3389/fmolb.2018.00093

10. Einstein, A. (1905). on the Movement of Small Particles Suspended in a Stationary Liquid Demanded By the Molecular-Kinetic Theory of Heat. Annalen Der Physik, 322(8), 549–560. https://doi.org/10.1002/andp.19053220806

11. Sutherland, W. (1905). LXXV. A dynamical theory of diffusion for non-electrolytes and the molecular mass of albumin. The London, Edinburgh, and Dublin Philosophical Magazine and Journal of Science, 9(54), 781–785. https://doi.org/10.1080/14786440509463331

12. Chandrasekhar, S. (1943). Stochastic problems in physics and astronomy. Reviews of Modern Physics, 15(1), 1–89. https://doi.org/10.1103/RevModPhys.15.1

13. Mueller, F., Mazza, D., Stasevich, T. J., & McNally, J. G. (2010). FRAP and kinetic modeling in the analysis of nuclear protein dynamics: What do we really know? In Current Opinion in Cell Biology (Vol. 22, Issue 3, pp. 403–411). https://doi.org/10.1016/j.ceb.2010.03.002

14. Day, C. A., & Kenworthy, A. K. (2009). Tracking microdomain dynamics in cell membranes. In Biochimica et Biophysica Acta - Biomembranes (Vol. 1788, Issue 1, pp. 245–253). https://doi.org/10.1016/j.bbamem.2008.10.024

15. He, L., & Niemeyer, B. (2003). A novel correlation for protein diffusion coefficients based on molecular weight and radius of gyration. Biotechnology Progress, 19(2), 544–548. https://doi.org/10.1021/bp0256059

16. He, L., & Niemeyer, B. (2003). A novel correlation for protein diffusion coefficients based on molecular weight and radius of gyration. Biotechnology Progress, 19(2), 544–548. https://doi.org/10.1021/bp0256059

17. Verkman, A. S. (2003). Diffusion in cells measured by fluorescence recovery after photobleaching. Methods in Enzymology, 360, 635–648. https://doi.org/10.1016/S0076-6879(03)60132-1

18. Lippincott-Schwartz, J., Altan-Bonnet, N., & Patterson, G. H. (2003). Photobleaching and photoactivation: Following protein dynamics in living cells. In Nature Reviews Molecular Cell Biology (Vol. 4, Issue SUPPL.).

19. Kitamura, A., Nakayama, Y., & Kinjo, M. (2015). Efficient and dynamic nuclear localization of green fluorescent protein via RNA binding. Biochemical and Biophysical Research Communications, 463(3), 401–406. https://doi.org/10.1016/j.bbrc.2015.05.084

20. Bacia, K., Kim, S. A., & Schwille, P. (2006). Fluorescence cross-correlation spectroscopy in living cells. Nature Methods, 3(2), 83–89. https://doi.org/10.1038/nmeth822

21. Dauty, E., & Verkman, A. S. (2004). Molecular crowding reduces to a similar extent the diffusion of small solutes and macromolecules: Measurement by fluorescence correlation spectroscopy. Journal of Molecular Recognition, 17(5), 441–447. https://doi.org/10.1002/jmr.709

22. Ramadurai, S., Holt, A., Krasnikov, V., Van Den Bogaart, G., Killian, J. A., & Poolman, B. (2009). Lateral diffusion of membrane proteins. Journal of the American Chemical Society, 131(35), 12650–12656. https://doi.org/10.1021/ja902853g

23. Cluzel, P., Surette, M., & Leibler, S. (2000). An ultrasensitive bacterial motor revealed by monitoring signaling proteins in single cells. Science, 287(5458), 1652–1655. https://doi.org/10.1126/science.287.5458.1652

24. Manley, S., Gillette, J. M., Patterson, G. H., Shroff, H., Hess, H. F., Betzig, E., & Lippincott-Schwartz, J. (2008). High-density mapping of single-molecule trajectories with photoactivated localization microscopy. Nature Methods, 5(2), 155–157. https://doi.org/10.1038/nmeth.1176

25. Digman, M. A., & Gratton, E. (2009). Analysis of diffusion and binding in cells using the RICS approach. Microscopy Research and Technique, 72(4), 323–332. https://doi.org/10.1002/jemt.20655

26. Waterman-Storer, C. M., Desai, A., Bulinski, J. C., & Salmon, E. D. (1998). Fluorescent speckle microscopy, a method to visualize the dynamics of protein assemblies in living cells. Current Biology, 8(22), 1227–1230. https://doi.org/10.1016/s0960-9822(07)00515-5

27. Gura Sadovsky, R., Brielle, S., Kaganovich, D., & England, J. L. (2017). Measurement of Rapid Protein Diffusion in the Cytoplasm by Photo-Converted Intensity Profile Expansion. Cell Reports, 18(11), 2795–2806. https://doi.org/10.1016/j.celrep.2017.02.063

28. Mika, J. T., & Poolman, B. (2011). Macromolecule diffusion and confinement in prokaryotic cells. In Current Opinion in Biotechnology (Vol. 22, Issue 1, pp. 117–126). https://doi.org/10.1016/j.copbio.2010.09.009

29. Miyawaki, A. (2011). Proteins on the move: Insights gained from fluorescent protein technologies. In Nature Reviews Molecular Cell Biology (Vol. 12, Issue 10, pp. 656–668). https://doi.org/10.1038/nrm3199

30. Kingsley, J. L., Bibeau, J. P., Mousavi, S. I., Unsal, C., Chen, Z., Huang, X., Vidali, L., & Tuzel, E. (2018). Understanding Boundary Effects and Confocal Optics Enables Quantitative FRAP Analysis in the Confined Geometries of Animal, Plant and Fungal Cells. Biophysical Journal, 114(3), 349a–350a. https://doi.org/10.1016/j.bpj.2017.11.1948

31. Ries, J., & Schwille, P. (2008). New concepts for fluorescence correlation spectroscopy on membranes. In Physical Chemistry Chemical Physics (Vol. 10, Issue 24, pp. 3487–3497). https://doi.org/10.1039/b718132a

32. Wachsmuth, M. (2014). Molecular diffusion and binding analyzed with FRAP. In Protoplasma (Vol. 251, Issue 2, pp. 373–382). https://doi.org/10.1007/s00709-013-0604-x

33. González-González, I. M., Jaskolski, F., Goldberg, Y., Ashby, M. C., & Henley, J. M. (2012). Measuring membrane protein dynamics in neurons using fluorescence recovery after photobleach. In Methods in Enzymology (Vol. 504, pp. 127–146). https://doi.org/10.1016/B978-0-12-391857-4.00006-9

34. Hardy, L. R. (2012). Fluorescence recovery after photobleaching (FRAP) with a focus on F-actin. Current Protocols in Neuroscience, SUPPL.61. https://doi.org/10.1002/0471142301.ns0217s61

35. Costantini, L., & Snapp, E. (2013). Probing endoplasmic reticulum dynamics using fluorescence imaging and photobleaching techniques. Current Protocols in Cell Biology, SUPPL.60. https://doi.org/10.1002/0471143030.cb2107s60

36. Zuleger, N., Kelly, D. A., & Schirmer, E. C. (2013). Considering discrete protein pools when measuring the dynamics of nuclear membrane proteins. Methods in Molecular Biology, 1042, 275–298. https://doi.org/10.1007/978-1-62703-526_2

37. Axelrod, D., Koppel, D. E., Schlessinger, J., Elson, E., & Webb, W. W. (1976). Mobility measurement by analysis of fluorescence photobleaching recovery kinetics. Biophysical Journal, 16(9), 1055–1069. https://doi.org/10.1016/S0006-3495(76)85755-4

38. Soumpasis, D. M. (1983). Theoretical analysis of fluorescence photobleaching recovery experiments. Biophysical Journal, 41(1), 95–97. https://doi.org/10.1016/S0006-3495(83)84410-5

39. Kang, M., Day, C. A., Drake, K., Kenworthy, A. K., & DiBenedetto, E. (2009). A generalization of theory for two-dimensional fluorescence recovery after photobleaching applicable to confocal laser scanning microscopes. Biophysical Journal, 97(5), 1501–1511. https://doi.org/10.1016/j.bpj.2009.06.017

40. Braga, J., Desterro, J. M. P., & Carmo-Fonseca, M. (2004). Intracellular macromolecular mobility measured by fluorescence recovery after photobleaching with confocal laser scanning microscopes. Molecular Biology of the Cell, 15(10), 4749–4760. https://doi.org/10.1091/mbc.E04-06-0496

41. Angelides, K. J., Elmer, L. W., Loftus, D., & Elson, E. (1988). Distribution and lateral mobility of voltage-dependent sodium channels in neurons. Journal of Cell Biology, 106(6), 1911–1925. https://doi.org/10.1083/jcb.106.6.1911

42. Seiffert, S., & Oppermann, W. (2005). Systematic evaluation of FRAP experiments performed in a confocal laser scanning microscope. Journal of Microscopy, 220(1), 20–30. https://doi.org/10.1111/j.1365-2818.2005.01512.x

43. Sprague, B. L., Pego, R. L., Stavreva, D. A., & McNally, J. G. (2004). Analysis of binding reactions by fluorescence recovery after photobleaching. Biophysical Journal, 86(6), 3473–3495. https://doi.org/10.1529/biophysj.103.026765

44. Deschout, H., Hagman, J., Fransson, S., Jonasson, J., Rudemo, M., Lorén, N., & Braeckmans, K. (2010). Straightforward FRAP for quantitative diffusion measurements with a laser scanning microscope. Optics Express, 18(22), 22886. https://doi.org/10.1364/oe.18.022886

45. Waharte, F., Steenkeste, K., Briandet, R., & Fontaine-Aupart, M. P. (2010). Diffusion measurements inside biofilms by image-based fluorescence recovery after photobleaching (FRAP) analysis with a commercial confocal laser scanning microscope. Applied and Environmental Microbiology, 76(17), 5860–5869. https://doi.org/10.1128/AEM.00754-10

46. Smisdom, N., Braeckmans, K., Deschout, H., vandeVen, M., Rigo, J.-M., De Smedt, S. C., & Ameloot, M. (2011). Fluorescence recovery after photobleaching on the confocal laser-scanning microscope: generalized model without restriction on the size of the photobleached disk. Journal of Biomedical Optics, 16(4), 046021. https://doi.org/10.1117/1.3569620

47. Braeckmans, K., Remaut, K., Vandenbroucke, R. E., Lucas, B., De Smedt, S. C., & Demeester, J. (2007). Line FRAP with the confocal laser scanning microscope for diffusion measurements in small regions of 3-D samples. Biophysical Journal, 92(6), 2172–2183. https://doi.org/10.1529/biophysj.106.099838

48. Stricker, J., Maddox, P., Salmon, E. D., & Erickson, H. P. (2002). Rapid assembly dynamics of the Escherichia coli FtsZ-ring demonstrated by fluorescence recovery after photobleaching. Proceedings of the National Academy of Sciences of the United States of America, 99(5), 3171–3175. https://doi.org/10.1073/pnas.052595099

49. Salmon, E. D., Leslie, R. J., Saxton, W. M., Karow, M. L., & McIntosh, J. R. (1984). Spindle microtubule dynamics in sea urchin embryos: Analysis using a fluorescein-labeled tubulin and measurements of fluorescence redistribution after laser photobleaching. Journal of Cell Biology, 99(6), 2165–2174. https://doi.org/10.1083/jcb.99.6.2165

50. Lajoie, P., Partridge, E. A., Guay, G., Goetz, J. G., Pawling, J., Lagana, A., Bharat, J., Dennis, J. W., & Nabi, I. R. (2007). Plasma membrane domain organization regulates EGFR signaling in tumor cells. Journal of Cell Biology, 179(2), 341–356. https://doi.org/10.1083/jcb.200611106

51. Zotter, A., Bäuerle, F., Dey, D., Kiss, V., & Schreiber, G. (2017). Quantifying enzyme activity in living cells. Journal of Biological Chemistry, 292(38), 15838–15848. https://doi.org/10.1074/jbc.M117.792119

52. Weiss, M. (2004). Challenges and artifacts in quantitative photobleaching experiments. Traffic, 5(9), 662–671. https://doi.org/10.1111/j.1600-0854.2004.00215.x

53. Pucadyil, T. J., & Chattopadhyay, A. (2006). Confocal fluorescence recovery after photobleaching of green fluorescent protein in solution. Journal of Fluorescence, 16(1), 87–94. https://doi.org/10.1007/s10895-005-0019-y

54. Kang, M., Day, C. A., Kenworthy, A. K., & DiBenedetto, E. (2012). Simplified equation to extract diffusion coefficients from confocal FRAP data. Traffic, 13(12), 1589–1600. https://doi.org/10.1111/tra.12008

55. Kitamura, A., & Kinjo, M. (2018). Determination of diffusion coefficients in live cells using fluorescence recovery after photobleaching with wide-field fluorescence microscopy. Biophysics and Physicobiology, 15(0), 1–7. https://doi.org/10.2142/biophysico.15.0_1

56. Gaigalas, A. K., Hubbard, J. B., McCurley, M., & Woo, S. (1992). Diffusion of bovine serum albumin in aqueous solutions. Journal of Physical Chemistry, 96(5), 2355–2359. https://doi.org/10.1021/j100184a063

57. Krouglova, T., Vercammen, J., & Engelborghs, Y. (2004). Correct diffusion coefficients of proteins in fluorescence correlation spectroscopy. Application to tubulin oligomers induced by Mg2+ and Paclitaxel. Biophysical Journal, 87(4), 2635–2646. https://doi.org/10.1529/biophysj.104.040717

58. Kuttner, Y. Y., Kozer, N., Segal, E., Schreiber, G., & Haran, G. (2005). Separating the contribution of translational and rotational diffusion to protein association. Journal of the American Chemical Society, 127(43), 15138–15144. https://doi.org/10.1021/ja053681c

59. Bertoni, M., Kiefer, F., Biasini, M., Bordoli, L., & Schwede, T. (2017). Modeling protein quaternary structure of homo- and hetero-oligomers beyond binary interactions by homology. Scientific Reports, 7(1). https://doi.org/10.1038/s41598-017-09654-8

60. Waterhouse, A., Bertoni, M., Bienert, S., Studer, G., Tauriello, G., Gumienny, R., Heer, F. T., De Beer, T. A. P., Rempfer, C., Bordoli, L., Lepore, R., & Schwede, T. (2018). SWISS-MODEL: Homology modelling of protein structures and complexes. Nucleic Acids Research, 46(W1), W296–W303. https://doi.org/10.1093/nar/gky427

61. Von Stetten, D., Noirclerc-Savoye, M., Goedhart, J., Gadella, T. W. J., & Royant, A. (2012). Structure of a fluorescent protein from Aequorea victoria bearing the obligate-monomer mutation A206K. Acta Crystallographica Section F: Structural Biology and Crystallization Communications, 68(8), 878–882. https://doi.org/10.1107/S1744309112028667

62. Hartinger, D., Heinl, S., Schwartz, H. E., Grabherr, R., Schatzmayr, G., Haltrich, D., & Moll, W. D. (2010). Enhancement of solubility in Escherichia coli and purification of an aminotransferase from Sphingopyxis sp. MTA144 for deamination of hydrolyzed fumonisin B1. Microbial Cell Factories, 9. https://doi.org/10.1186/1475-2859-9-62

63. Hartinger, D., Heinl, S., Schwartz, H. E., Grabherr, R., Schatzmayr, G., Haltrich, D., & Moll, W. D. (2010). Enhancement of solubility in Escherichia coli and purification of an aminotransferase from Sphingopyxis sp. MTA144 for deamination of hydrolyzed fumonisin B1. Microbial Cell Factories, 9. https://doi.org/10.1186/1475-2859-9-62

64. Kumar, M., Mommer, M. S., & Sourjik, V. (2010). Mobility of cytoplasmic, membrane, and DNA-binding proteins in Escherichia coli. Biophysical Journal, 98(4), 552–559. https://doi.org/10.1016/j.bpj.2009.11.002

65. Mika, J. T., Schavemaker, P. E., Krasnikov, V., & Poolman, B. (2014). Impact of osmotic stress on protein diffusion in Lactococcus lactis. Molecular Microbiology. https://doi.org/10.1111/mmi.12800

66. Konopka, M. C., Sochacki, K. A., Bratton, B. P., Shkel, I. A., Record, M. T., & Weisshaar, J. C. (2009). Cytoplasmic protein mobility in osmotically stressed Escherichia coli. Journal of Bacteriology, 91(1), 231–237. https://doi.org/10.1128/JB.00536-08

67. Sochacki, K. A., Shkel, I. A., Record, M. T., & Weisshaar, J. C. (2011). Protein diffusion in the periplasm of E. coli under osmotic stress. Biophysical Journal, 100(1), 22–31. https://doi.org/10.1016/j.bpj.2010.11.044

68. Van Den Bogaart, G., Hermans, N., Krasnikov, V., & Poolman, B. (2007). Protein mobility and diffusive barriers in Escherichia coli: Consequences of osmotic stress. Molecular Microbiology, 64(3), 858–871. https://doi.org/10.1111/j.1365-2958.2007.05705.x

